# Time-resolved structured illumination microscopy reveals key principles of Xist RNA spreading

**DOI:** 10.1101/2020.11.24.396473

**Authors:** Lisa Rodermund, Heather Coker, Roel Oldenkamp, Guifeng Wei, Joseph Bowness, Bramman Rajkumar, Tatyana Nesterova, David Pinto, Lothar Schermelleh, Neil Brockdorff

**Affiliations:** Department of Biochemistry, University of Oxford, South Parks Rd, Oxford, OX1 3QU, UK; Division of Gene Regulation, Netherlands Cancer Institute, Plesmanlaan 121, Amsterdam, 1066 CX, Netherlands

## Abstract

Xist RNA directs the process of X-chromosome inactivation in mammals by spreading *in cis* along the chromosome from which it is transcribed and recruiting chromatin modifiers to silence gene transcription. To elucidate mechanisms of Xist RNA *cis*-confinement, we established a sequential dual color labeling, super-resolution imaging approach to trace individual Xist RNA molecules over time, enabling us to define fundamental parameters of spreading. We demonstrate a feedback mechanism linking Xist RNA synthesis and degradation, and an unexpected physical coupling between preceding and newly synthesized Xist RNA molecules. Additionally, we show that the protein SPEN, a key factor for Xist-mediated gene-silencing, has a distinct function in Xist RNA localization, stability and in coupling behavior. Our results provide important insights towards understanding the unique dynamic properties of Xist RNA.

**One Sentence Summary:** Visualizing Xist RNA dynamics in single cells during X chromosome inactivation

## Introduction

X chromosome inactivation (XCI) is the mechanism that evolved in mammals to equalize levels of X-linked gene expression in XX females relative to XY males [reviewed in (*1*)]. The XCI process is controlled by a 17 kb long non-coding RNA, Xist (X inactive specific transcript) that functions *in cis* to silence the chromosome from which it is transcribed. Xist RNA localizes *in cis* across the length of the inactive X chromosome (Xi) elect, recruiting silencing factors that mediate chromatin modification and transcriptional inactivation of underlying genes [reviewed in (*2*)]. Expression of *Xist* transgenes on autosomes also results in chromosome silencing *in cis* (*3,4*).

A key challenge in X inactivation research is to understand the basis for *cis*-limited localization of Xist RNA. Early studies revealed that Xist RNA concentrates over chromosomal regions that have a high gene density (*5*), and that Xist RNA interacts tightly with the nuclear matrix fraction (*6*). Consistent with the latter observation, the nuclear matrix proteins hnRNPU and Ciz-1 are required to anchor Xist RNA to the Xi territory (*7–10*). Related to these findings, super-resolution 3D-SIM imaging has revealed that Xist RNA molecules reside in distinct foci adjacent to hnRNPU in interchromatin channels that pervade the Xi territory (*11*). More recently, mapping of chromatin sites associated with Xist RNA using RNA antisense purification (RAP) (*12*), have revealed initial preferred localization to regions that are in close 3D proximity to the site of transcription, with subsequent spreading across the chromosome (*13*). These findings support that preferred chromatin sites for Xist RNA association are not defined by underlying DNA sequence.

Despite the aforementioned progress, our understanding of *cis*-limited localization of Xist RNA remains rudimentary. We recently proposed a model that invokes that the range over which Xist RNA spreads is a function of Xist RNA abundance, based on synthesis and degradation rates, and the dynamics of the anchoring interaction of Xist ribonucleoprotein particles (RNPs) with the nuclear matrix (*14*). To test this hypothesis, we set out to quantify the dynamic behavior of Xist RNPs. Previous studies have described approaches to fluorescently tag Xist RNA in living cells using fluorescent proteins (*15,16*), but these systems are amenable only to conventional diffraction-limited fluorescence microscopy, meaning it is not possible to observe individual Xist RNPs. To overcome this limitation, we developed a novel methodology, RNA-SPLIT (<u>S</u>equential <u>P</u>ulse <u>L</u>ocalization <u>I</u>maging over <u>T</u>ime), that allows measurement of Xist RNP dynamics and localization using super-resolution three-dimensional structured illumination microscopy (3D-SIM). We use this approach to define critical parameters underpinning the behavior of Xist RNPs, and to show a novel role for the silencing protein SPEN in Xist RNA behavior.

## Development of RNA-SPLIT to analyze Xist RNA dynamics

We set out to measure the dynamic behavior of individual Xist RNA molecules using superresolution 3D-SIM. The Bgl-stem-loop system (*17*) was employed for efficient labelling of inducible Xist RNA from its endogenous location on the X chromosome (Chr X) or from an autosomal Xist transgene integration site on chromosome 15 (Chr 15) (Fig. 1A). In combination with HaloTag technology, photostability and brightness could be increased sufficiently for the application of 3D-SIM, allowing the detection of distinct Xist RNPs within sub-nuclear domains, referred to henceforth as Xist clusters (Fig. 1, A and B). We inferred that focal signals seen with HaloTag staining correspond to individual Xist RNPs based on the evidence of HaloTag foci having very similar intensities (see methods), and a striking co-localization with focal signals of immunofluorescence-detected Xist RNA binding protein Ciz-1 (fig. S1A). Insertion of the Bgl-stem-loop array had no effect on Xist-mediated silencing, determined using allelic chromatin RNA sequencing (ChrRNA-seq) (fig. S1B), and tagged Xist RNA was shown to recruit known co-factors and to establish Xi-specific histone modifications (fig. S1C). Similar results were obtained for mESCs expressing the Chr 15 Xist transgene (fig. S1, D and E). Both cell lines showed an induction efficiency of over 80 % with no detectable promoter leakiness (fig. S1, F and G).

**Fig. 1.**
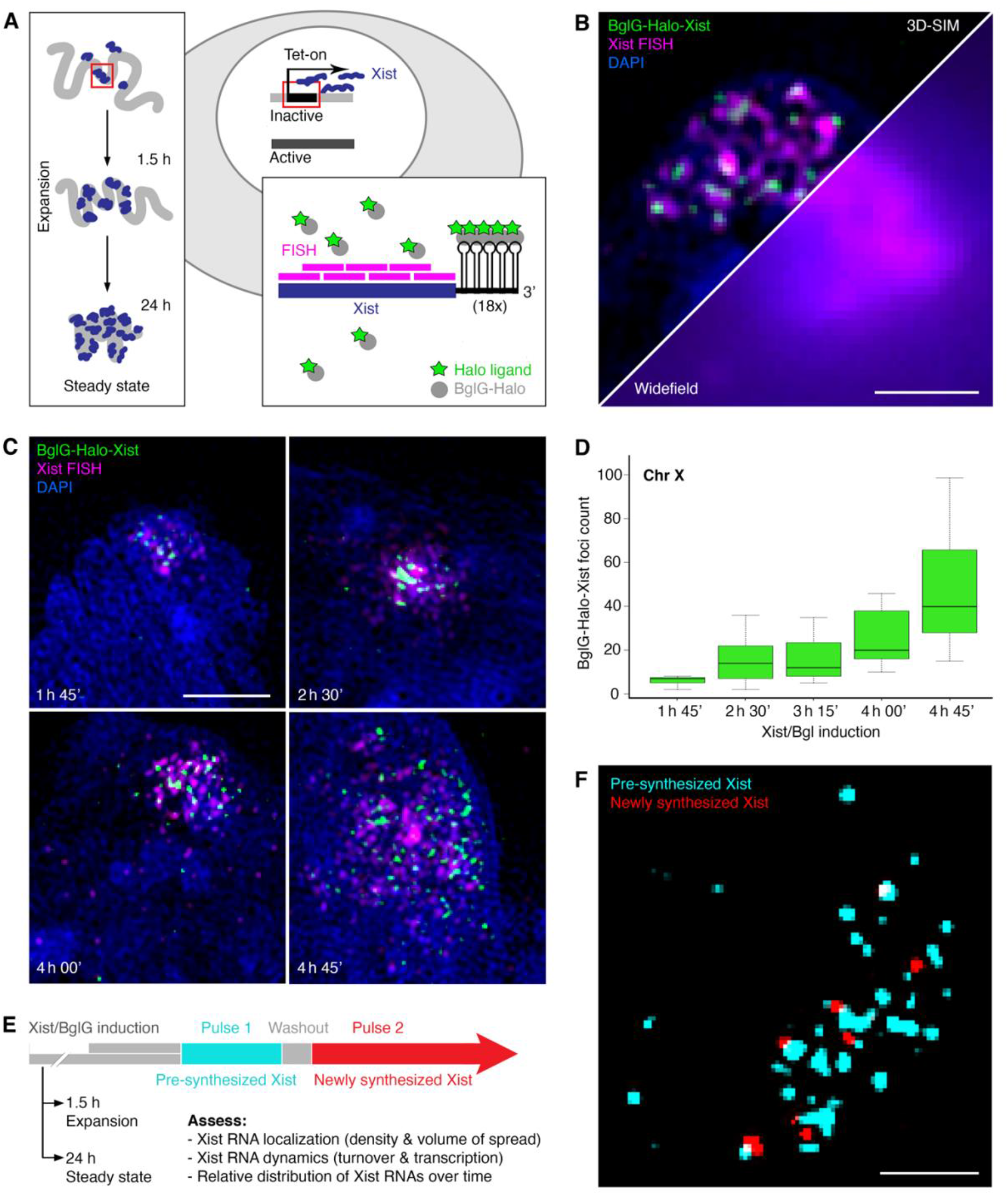
Measuring Xist RNA behavior with RNA-SPLIT. (**A**) Schematic illustrating HaloTag labelling of Xist RNA in mESCs via the BglG/Bgl stem loop system. (**B**) Combined Xist RNA FISH with HaloTag staining of BglG-Halo-Xist visualized by 3D-SIM and conventional widefield microscopy for comparison. DNA is counterstained with DAPI. Single cross-section of a 3D image stack displayed. Scale bar: 1 μm. (**C**) Representative 3D-SIM images (z-projections) of Xist clusters labelled by RNA FISH and HaloTag staining at different time points of Xist induction. Scale bar: 2 μm. (**D**) Boxplots showing average Xist foci numbers after Xist induction for different times determined from 3D-SIM analysis of Halo-BglG labelled Xist RNPs. n = 11 cells/time point. (**E**) Schematic illustrating the RNA-SPLIT regimen and the parameters of Xist RNA behavior measured in this study. (**F**) Representative 3D-SIM image (single cross section) of Xist RNA-SPLIT showing newly synthesized (red) and pre-synthesized (cyan) Xist RNA molecules. Scale bar: 1 μm.

We identified a time window from 1.5 h until 5 h post-induction during which we could quantify a gradual increase of Xist RNPs within the volume of the Xist cluster (Fig. 1, C and D), referred to henceforth as expansion phase. A later timepoint (24 h) was taken as representative of Xist RNA behavior at steady state. We applied 3D-SIM live cell imaging to detect distinct Xist foci over a period of 5 min, but found that the foci were largely static, even at the fastest possible acquisition rates (movie S1). An alternative approach to track Xist RNA molecules using single-particle tracking after photoactivation (PALM-SPT) was unsuccessful due to the high background signal from non-RNA bound BglG-Halo fusion protein. To overcome these limitations, we developed RNA-SPLIT, in which successive rounds of labelling with different HaloTag ligands performed in live cells allowed separate imaging of the first and subsequent rounds of Xist RNA synthesis (henceforth referred to as pre-synthesized and newly synthesized Xist RNA respectively), analyzed with super-resolution 3D-SIM after formaldehyde fixation (Fig. 1, E and F). Varying labelling times in combination with a custom image processing and quantitative analysis pipeline enabled measurement of various aspects of RNA dynamics and localization, notably density, spreading behavior, RNA turnover and transcription dynamics (Fig. 1E and fig. S2, A and B).

## Chromatin environment and rate of transcription modulate Xist RNA stability

We applied the RNA-SPLIT protocol to assess Xist RNA turnover following induction during expansion phase (after 1.5 h) or steady state (after 24 h), both on Chr X and Chr 15. A gradual reduction of pre-synthesized RNA was observed over a 4 h time course, with higher stability seen for Chr X compared to Chr 15 expressed Xist RNA (Fig. 2A-C). In both cell lines, stability was higher in steady state compared to expansion phases. The latter observation indicates that Xist RNA stability increases with XCI progression independent of chromosomal context. We noted that turnover of pre-synthesized RNA is more evident towards the periphery of Xist RNP clusters (Fig. 2B). Results for Xist RNA turnover rates were validated using a different method, SLAM-seq (*18*) (Fig. 2D and fig. S3, A-D). The use of SLAM-seq further allowed us to ascertain that insertion of the Bgl-stem-loop array does not significantly alter Xist transcript stability (fig. S3, A-D).

**Fig. 2.**
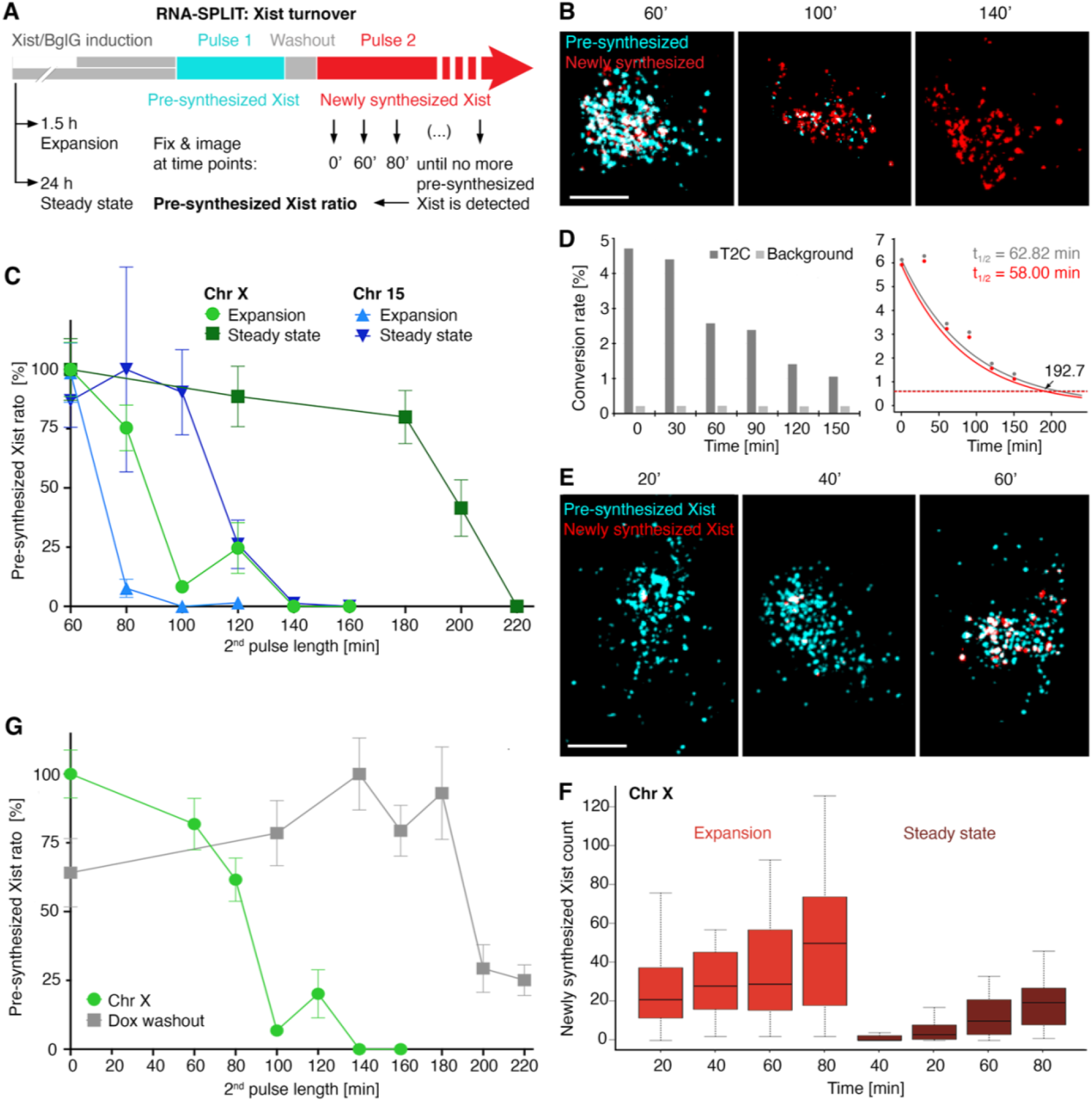
Xist RNA stability is defined by its chromatin environment and rate of transcription. (**A**) Schematic illustrating RNA-SPLIT regimen to assess Xist RNA turnover. (**B**) Representative 3D-SIM images (z-projections) of RNA-SPLIT experiment to assess Xist RNP turnover. Note pre-synthesized Xist RNPs turnover more rapidly at the periphery of the cluster. Scale bar: 2 μm. (**C**) Plot showing quantification of Chr X and Chr 15 Xist RNA turnover during expansion and steady state. n = 20 cells/time point. (**D**) Plots of SLAM-seq experiments showing decrease of T→C conversions over time (left) and data for Xist RNA (BglG-Halo tagged) replicates fitted to exponential decay curves (right) from which t1/2 values are derived. The grey and red curves represent the raw T to C conversion and background-corrected conversion rates respectively. The red dashed line indicated 10% of original signal. (**E**) Representative 3D-SIM images (z-projections) for Chr X expressed Xist RNA during expansion. Scale bar: 2 μm. (**F**) Boxplots showing Xist RNA transcription over time during expansion and steady state phases. n = 20 cells/time point. (**G**) Plot showing effect of Dox washout on Xist RNP turnover. n = 20 cells/time point.

Quantitation of the average number of Xist RNPs in cells indicated a general decrease from expansion to steady state phase (fig. S3E), apparently at odds with the observed increase in stability. Further analysis revealed that this is accounted for by significantly elevated Xist transcription rates in expansion compared with steady state, as determined using RNA-SPLIT analysis of both Chr X and Chr 15 expressed Xist RNA (Fig. 2, E and F, and fig. S3F). We noted increased transcription rates of Xist expressed in the Chr 15 compared to Chr X context. These observations suggest that the abundance of Xist RNA may be regulated by a feedback mechanism that balances rates of synthesis and degradation.

A previous study reported increased Xist RNA stability following inhibition of transcription by actinomycin D (*15*), also pointing to a link between Xist RNA synthesis and turnover. We re-examined this finding using doxycycline washout to terminate Xist expression in preference to actinomycin D treatment which can lead to confounding indirect effects. Doxycycline washout performed 1.5 h post-induction (fig. S3G) resulted in a dramatic increase in Xist RNA stability (Fig. 2G). This result further supports the existence of a promoter-independent feedback mechanism that links rates of Xist RNA transcription and turnover.

## Time-resolved analysis of Xist RNA localization

We applied RNA-SPLIT to obtain temporally resolved 3D localization information at the level of individual Xist RNPs in single cells. The site of Xist transcription was approximated based on the density and fluorescence intensity of newly synthesized Xist RNA signal. Using this approach, we found that the transcription site is on average located centrally within Xist RNA clusters in expansion phase and slightly more peripherally at steady state (fig. S4, A and B). We went on to measure 3D distances of newly synthesized and pre-synthesized Xist RNA molecules from the transcription site (Fig. 3A). By dividing the Xist RNA cluster into three concentric zones, we were able to quantify the relative distribution of Xist molecules with respect to distance from the transcription site over time (Fig. 3B). This analysis revealed that newly synthesized but not pre-synthesized Xist RNPs spread in 3D from the transcription site towards the periphery of the Xist cluster over a 1 h time course during expansion phase. We calculated the average expansion rate to be ~8 μm^3^/h. Taking into account Xist RNA-seq data collected over a similar time course (*13*), and also H2AK119ub1 ChIP-seq analysis shown below, we infer that the observed spreading reflects Xist RNPs being transported initially to sites in close proximity to the Xist locus (see also discussion). 3D spread was not measurable using the same time course during steady state (Fig. 3B). Possible explanations include differences in Xist RNA turnover/transcription rate, or a more stochastic spread of Xist RNA as was proposed previously based on RAP-seq and CHART-seq experiments performed at later post-induction time points (*13,19*).

**Fig. 3.**
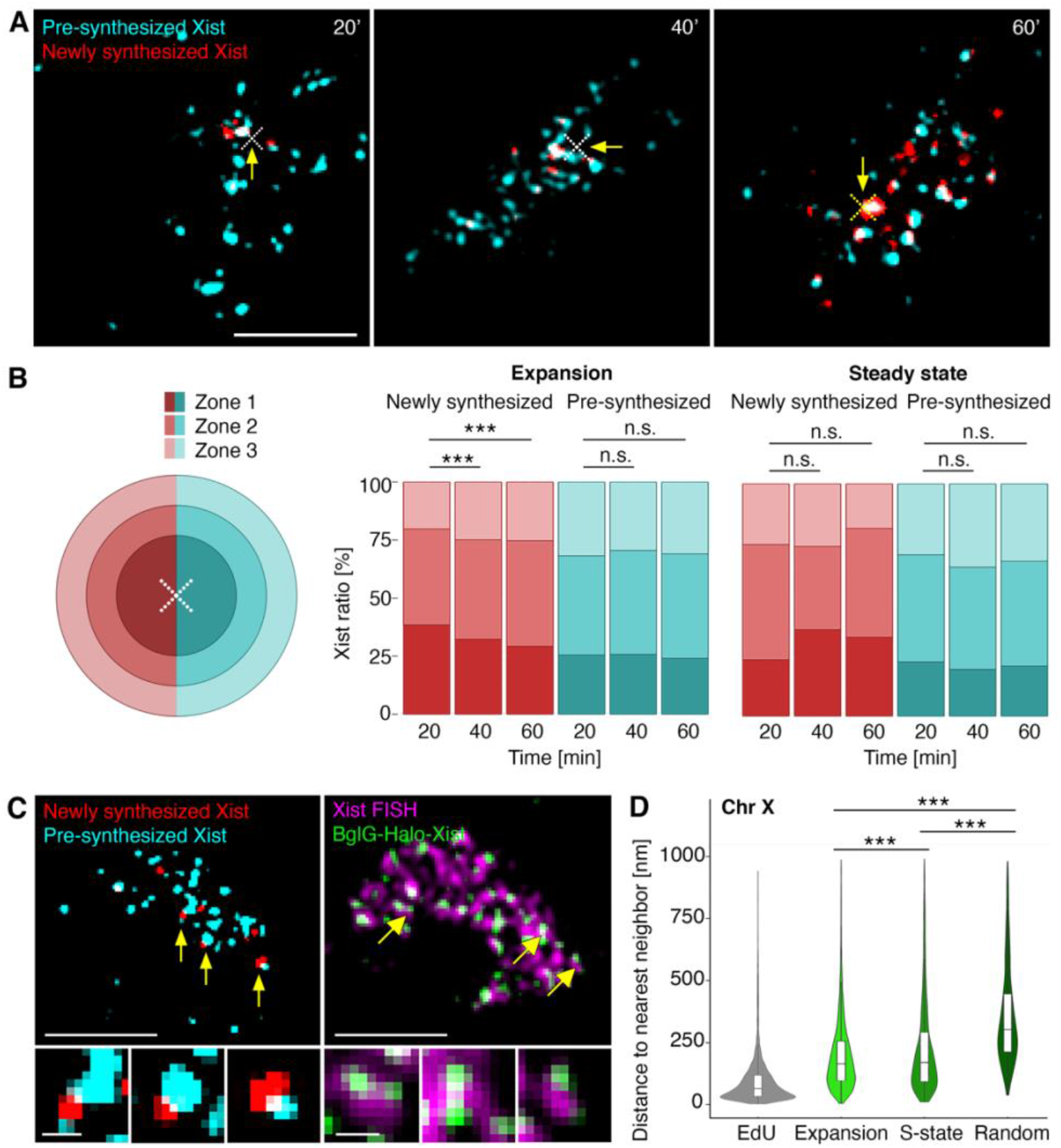
Xist RNP localization behavior. (**A**) Representative 3D-SIM images (single z-section) of RNA-SPLIT illustrating Xist RNP spread in expansion phase. Approximated Xist TSS is indicated with white cross. Scale bar: 2 μm. (**B**) Bar chart quantifying range of newly synthesized and pre-synthesized Xist RNPs during expansion and steady state phases. Each signal was assigned to a zone as illustrated in schematic (right). n = 40 cells/time point. Significance determined by unpaired two-sample Wilcoxon test. (**C**) Representative 3D-SIM image (single z-section) of RNA-SPLIT (left) and Xist RNA FISH combined with BglG HaloTag labelling (right) illustrating coupling of pre-synthesized and newly synthesized Xist RNPs during expansion phase. Lower panels show examples of coupled Xist. Single cross-sections. Selected couplets (arrows) are expanded in panels below. Scale bars: 2 μm (main image) or 200 nm (lower panels). (**D**) Violin plots quantifying Xist RNP coupling in expansion and steady state phases. n = 200 cells/time point. Colocalization offset is defined using EdU control. Significance determined by unpaired two-sample Wilcoxon test.

Interestingly, RNA-SPLIT analysis revealed the frequent occurrence of pairwise association of pre-synthesized and newly synthesized (coupled) Xist RNP signals (Fig. 3C, left panel). This phenomenon was also seen in experiments combining Xist RNA FISH with HaloTag staining, where closely coupled HaloTag signals are seen to closely associate with larger Xist RNA FISH foci (Fig. 3C, right panel). The possibility that this effect is due to detection of Xist RNPs double-labeled with the different HaloTag ligands was ruled out using a dual-color control (see methods) (fig. S5A).

We used nearest neighbor analysis (NNA) of 3D-SIM RNA-SPLIT images to quantify Xist RNP coupling at expansion and steady state in both Chr X and Chr 15 mESCs. Estimated median distances were in the range 160-180 nm (Fig. 3D and fig. S5B). To determine the technical offset for colocalizing nuclear focal signals, we used 5-ethynyl-2’-deoxyuridine (EdU) pulse replication labelled cells that were simultaneously detected with two colors (*20*). Taking into account the minimum resolvable separation using 3D-SIM of ~60 nm obtained for the NNA of the dual-color EdU control sample (Fig. 3D), and an estimated circumference of Xist RNPs of 50 nm (derived from the ratio of Xist RNA and rRNA length and the known size of a ribosome), these observations demonstrate very close spatial association of individual pre-synthesized and newly synthesized Xist RNPs. This finding is further emphasized by comparison with a randomized NNA control (Fig. 3D and fig. S5B). Importantly, there is no equivalent frequency of coupling of pre-synthesized with pre-synthesized, or newly synthesized with newly synthesized Xist RNPs, indicating that Xist RNPs are not synthesized/transported as couplets but rather come together at distant bound sites.

## Application of RNA-SPLIT to investigate the role of Ciz-1 in Xist RNA behavior

To gain further insight into the mechanistic aspects of Xist RNA behavior we applied RNA-SPLIT while perturbing factors known to be involved in *cis*-limited Xist RNA localization. Firstly, we generated a knockout (KO) of the gene encoding Ciz-1 (fig. S6, A-D), a nuclear matrix protein which interacts with Xist RNA and plays a pivotal role in anchoring Xist RNPs in somatic cells (*9,10*). In line with previous analyses (*9,10*), we observed little or no effects on Xist RNA localization in undifferentiated mESCs (Fig. S6E and fig. S6F). Accordingly, Xist transcript stability, transcription dynamics and coupling were largely unaffected and moreover, Xist-mediated silencing determined using ChrRNA-seq was unaltered (fig. S7, A-E). In marked contrast, in differentiated neural precursor cells (NPCs) derived from the Ciz-1 KO mESCs we observed a dramatic dispersal of Xist RNPs throughout the entire nucleus (Fig. 4, A and B). We quantified the localization defect in NPCs by comparing inter-molecule distances in Ciz-1 KO and WT NPCs (Fig. 4C). In WT NPCs the density of Xist RNPs increased significantly as differentiation progressed, whilst in Ciz-1 KO NPCs density progressively decreased, (Fig. 4C).

**Fig. 4.**
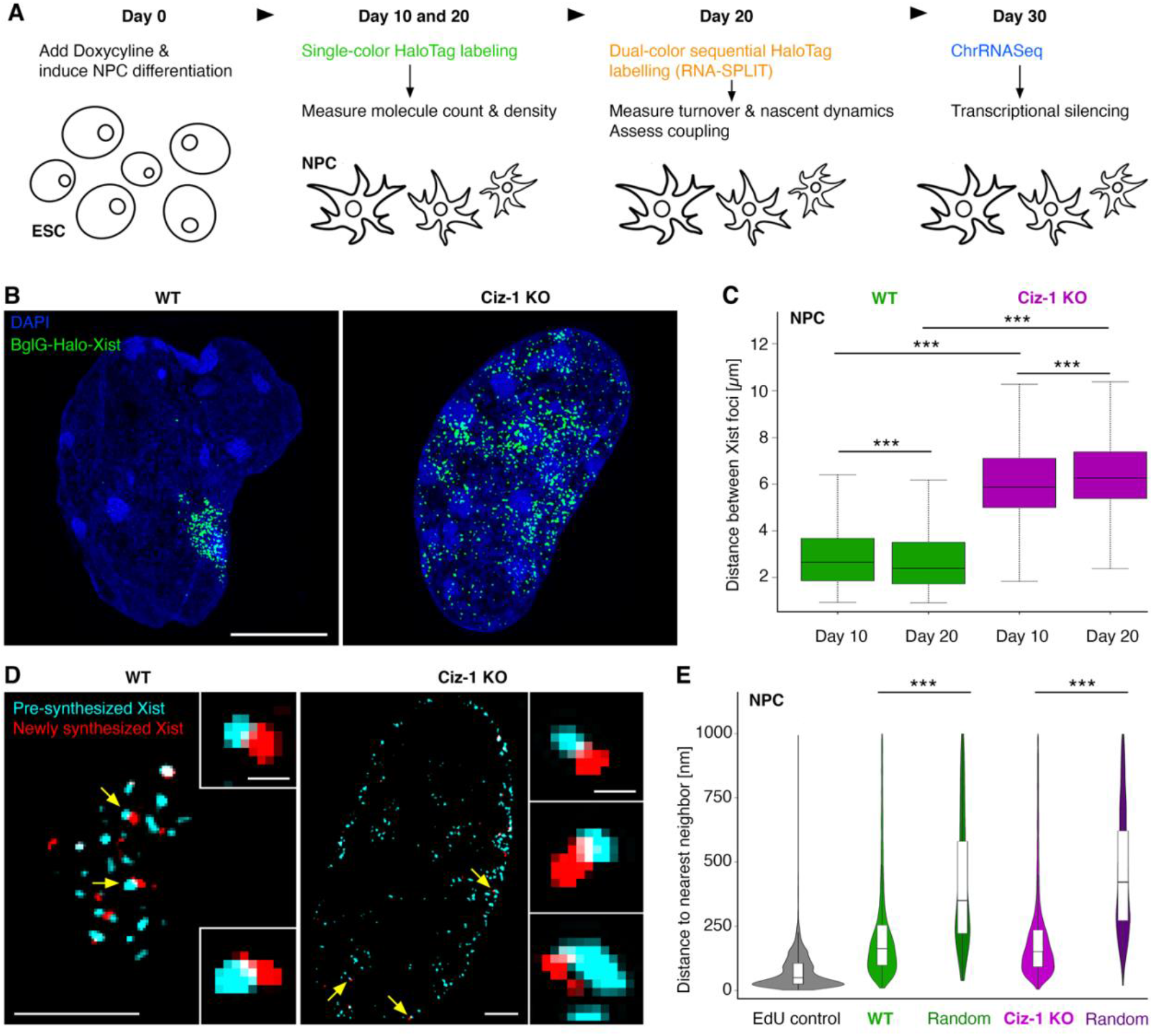
Investigation of Ciz-1 function in regulating Xist RNA behavior. (**A**) Schematic illustrating differentiation of WT and Ciz-1 KO ESCs into NPCs. (**B**) Representative 3D-SIM images (z-projections) of Xist RNPs in NPCs at day 10 of differentiation. Scale bar: 2 μm. (**C**) Boxplots representing Xist RNP density in Ciz-1 KO NPCs. n = 40 cells/time point. Significance determined using unpaired two-sample Wilcoxon test. (**D**) Representative 3D-SIM images (single z-sections) of RNA SPLIT experiments for WT and Ciz-1 KO NPCs after 20 days of differentiation showing coupling of pre-synthesized and newly synthesized Xist RNPs. Selected couplets (arrows) are expanded in panels to the right. Scale bars: 2 μm (main images), and 200 nm (expanded images). (**E**) Corresponding nearest neighbor analysis between Xist RNP species in WT and Ciz-1 KO NPCs. n = 150 cells/time point. Significance determined using unpaired two-sample Wilcoxon test.

Despite the observed delocalization of Xist RNA, ChrRNA-Seq analysis demonstrated that there was no major difference in transcriptional silencing of the Xi (fig. S8A), consistent with the fact that Ciz-1 KO female mice are viable (*9*). Both RNA-SPLIT analysis and ChrRNA-Seq demonstrated an increased Xist RNA molecule count in NPCs as compared to mESCs, with an even more dramatic increase in Ciz-1 KO NPCs (fig. S8, B and C). Further analysis by RNA-SPLIT revealed that this is linked to increased Xist transcription, most notably in Ciz-1 KO NPCs (fig. S8D). Additionally, Xist RNA stability is increased in NPCs compared to mESCs (fig. S8E). These observations indicate that the feedback between Xist transcription and turnover is disrupted in Ciz-1 KO NPCs.

We went on to apply RNA-SPLIT to assess coupling of pre-synthesized and newly synthesized Xist RNPs in NPCs. Coupling was readily observable in WT NPCs, demonstrating that the phenomenon is common to both undifferentiated and differentiated cells types (Fig. 4D). Strikingly, we also observed coupling in Ciz-1 KO NPCs, despite dispersal of Xist RNPs throughout the nucleus (Fig. 4E). This finding demonstrates that coupling of newly synthesized and pre-synthesized Xist RNPs is independent of cis-limited localization to the X chromosome.

## A role for SPEN in Xist RNA localization

The protein SPEN, which binds to the A-repeat region of Xist RNA via a triple RRM domain, plays a central role in Xist-mediated gene silencing (*8,17,21,22*). Silencing activity has been linked to a C-terminal SPOC domain that interacts with the NCoR-HDAC3 co-repressor complex (*23–25*). In recent work we reported atypical Xist RNA clusters and reduced levels of Xist RNA resulting from SPEN loss of function (RRM deletion), and similarly following deletion of the Xist A-repeat (*26*). Consistent with these observations, prior work reported that the A-repeat is critical for *in cis* localization of a truncated Xist RNA transgene (*27*). To further investigate a possible role for SPEN in Xist RNA localization we engineered the SPEN RRM deletion in mESCs with endogenous inducible BglG-Halo-Xist (Fig. 5A and fig. S9A). Analysis by ChrRNA-seq confirmed a major silencing defect, (Fig. S9B), consistent with prior studies (*25,26*). 3D-SIM combined with HaloTag staining revealed a visible defect in Xist RNA localization, both in expansion and steady state phases (Fig. 5B and fig. S9C).

**Fig. 5.**
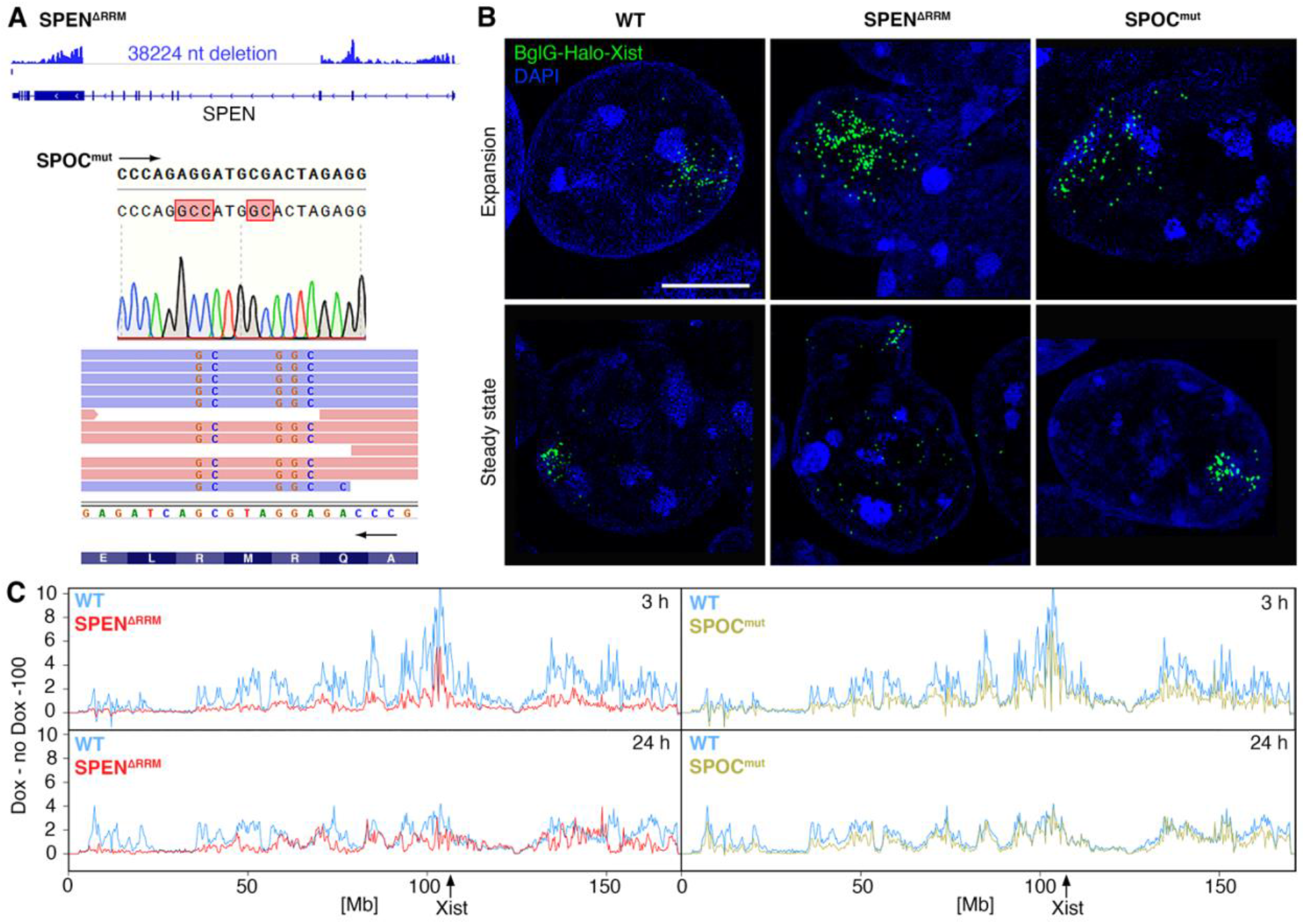
A role for SPEN in Xist RNA localization. (**A**) UCSC genome browser screenshot with ChrRNA-seq track confirming 38224 nucleotide deletion of sequences encoding the SPEN RRM region (top), with indication of nucleotide substitutions in the SPOC domain mutant confirmed by Sanger sequencing (bottom). (**B**) Representative 3D-SIM images (z-projections) showing Xist RNP clusters in WT and SPEN mutant mESCs as indicated in both expansion and steady state phases. Scale bar: 2 μm. (**C**) Plots showing gain of H2AK119ub1 within 250 kb windows on chromosome X determined by ChIP-seq analysis in SPEN RRM del cells (left) and SPOC domain mutant cells (right) after 3 h (top) and 24 h (bottom) induction with doxycycline. The position of the Xist locus is indicated (arrow).

Prior studies have determined a close correlation of Xist-dependent Polycomb histone modifications and Xist RNA binding as determined by RAP (*13*), and we therefore carried out ChIP-seq analysis of Polycomb-mediated H2AK119ub1 to approximate the chromosomal distribution of Xist RNA in SPEN RRM deletion mESCs. After induction of Xist for 3 h or 24 h, representing expansion and steady state, respectively, we observed a marked reduction in H2AK119ub1, in particular at sites further away from the *Xist* locus (Fig. 5C and fig. S9D). This observation indicates that SPEN is important for the long-range spread of Xist RNA.

Aberrant Xist RNA localization in SPEN null mESCs may be a consequence of abrogated gene silencing, or alternatively may represent a distinct function of SPEN. To discriminate these possibilities, we generated mESCs with mutations in the SPOC domain previously shown to disrupt interaction with the NCoR/HDAC3 co-repressor (*28*) (Fig. 5A). ChrRNA-seq analysis revealed that mutation of the SPEN SPOC domain abrogates Xist-mediated gene silencing, but to a lesser degree compared with the SPEN RRM deletion (fig. S9B). This observation is consistent with a recent study that analyzed a SPOC domain deletion mutant (*25*). Interestingly, aberrant localization of Xist RNA was not apparent in the SPEN SPOC mutation mESCs (Fig. 5B), and moreover, there was little or no effect on Xist RNA distribution as assessed by ChIP-seq analysis of H2AK119ub1 (Fig. 5C and fig. S9D). These observations suggest a SPOC independent function of SPEN in X inactivation that is linked to a role in Xist RNA localization.

## SPEN modulates Xist RNA dynamics and is required for coupling Xist RNPs

We went on to quantify the effect of SPEN mutations on Xist RNA behavior. The SPEN RRM deletion (and to a lesser extent mutation of the SPOC domain), resulted in a small decrease in the efficiency of Xist RNA induction (Fig. S9C). While Xist RNP density was significantly decreased in both cell lines, the volume of spread was more affected in the SPEN RRM deletion (Fig. 6A and fig. S10A). Here, Xist occupied on average 70 % of the nucleus in contrast to around 15 % in the WT (Fig. 6B). The SPEN RRM deletion (but not the SPOC domain mutant) results in reduced levels of Xist RNA at steady state (fig. S10, B and C). This finding was not attributable to decreased Xist transcription, which was actually significantly increased both in expansion and steady state (Fig. 6C and fig. S10D), but rather to a significant decrease in Xist RNA stability (Fig. 6D).

**Fig. 6.**
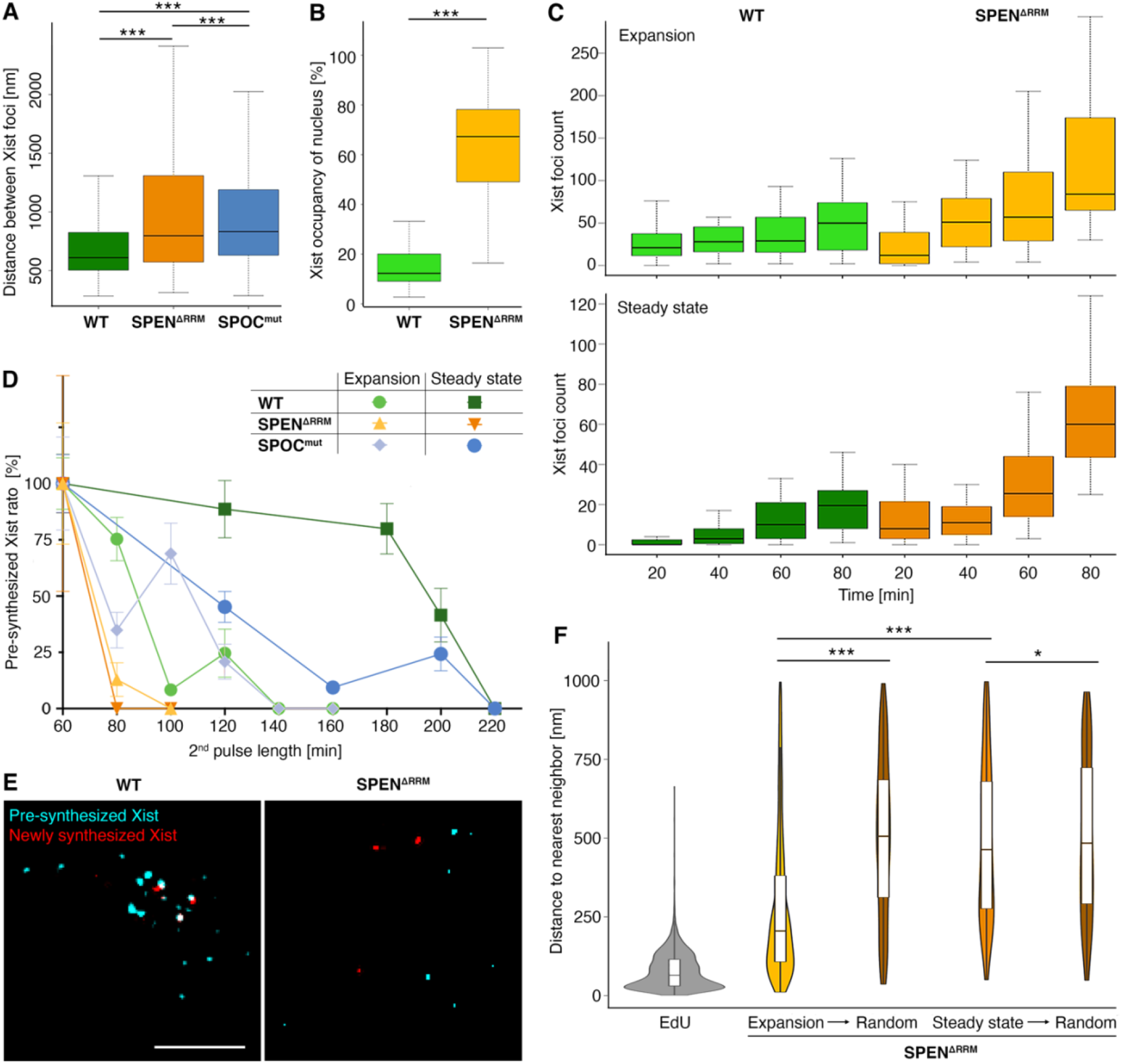
SPEN modulates Xist RNP stability and coupling. (**A**) Boxplots showing quantification of Xist RNP density in SPEN RRM del and SPOC domain mutant cells. n = 240 cells/cell line. Significance determined using unpaired two-sample Wilcoxon test. (**B**) Boxplots showing Xist cluster volume as proportion of nuclear volume in SPEN RRM deletion cells. n = 120 cells/cell line. Significance determined using unpaired two-sample Wilcoxon test. (**C**) Boxplots illustrating Xist RNA transcription in SPEN RRM del during expansion and steady state. n = 20 cells/ time point. (**D**) Plot showing Xist RNP turnover in mESCs as indicated during expansion (exp) and steady state (ss). n = 20 cells/time point. (**E**) Representative 3D-SIM images (single z-section) of RNA-SPLIT illustrating abrogation of Xist RNP coupling in SPEN RRM del mESCs. Scale bar: 2 μm. (**F**) Violin plots showing abrogated Xist RNP coupling in SPEN RRM del mESCs. n = 200 cells/time point. Significance determined using unpaired two-sample Wilcoxon test.

In a final series of experiments, we applied RNA-SPLIT to determine if SPEN mutations affect Xist RNP coupling. While mutation of the SPOC domain had no apparent effect, deletion of the SPEN RRM strongly abrogated coupling (Fig. 6, E and F, and fig. S10, E and F). Loss of coupling was evident both in expansion and steady state phases, although most dramatically in the latter. Together these results demonstrate a previously unappreciated role for SPEN in Xist RNA behavior that is independent of gene-silencing functions mediated by the SPOC domain, and likely involves regulation of Xist RNP stability, localization and/or coupling.

## Discussion

In this work we identify two novel parameters that need to be considered in modelling Xist RNA behavior. First, we find that Xist RNP abundance during expansion and steady state is maintained by a feedback mechanism that links rates of synthesis and degradation. Factors regulating the native Xist promoter are unlikely to be involved as our experiments analyzed Xist RNA expressed from the heterotypic doxycycline promoter. The feedback mechanism is unaffected by mutation of the SPEN SPOC domain, indicating that it does not require Xist-mediated gene silencing, but is perturbed upon SPEN RRM deletion, and in Ciz-1 KO NPCs. Accordingly, we suggest that anchoring of Xist RNPs to the nuclear matrix, which is perturbed in SPEN RRM deletion mESCs and Ciz-1 KO NPCs, may be important for linking rates of transcription and turnover. Second, we find a remarkable spatial co-localization of individual newly synthesized with pre-synthesized Xist RNPs that we refer to as coupling. The spatial and temporal information we obtained indicates that coupling takes place at Xist RNP binding sites rather than at the Xist transcription site. We speculate that a conformational switch occurs when Xist RNPs anchor to the nuclear matrix, and that this facilitates coupling, either with a mobile/diffusing Xist RNP, or a second anchored Xist RNP located at a different site, but brought into proximity by dynamic movement of the chromosome (fig. S11, A and B).

Using RNA-SPLIT we determined that during the expansion phase, Xist RNA clusters enlarge radially at a rate of ~8 μm^3^/h. The behavior that we observe is consistent with the proposal that Xist RNA initially spreads to sites that are proximal in 3D, and then distributes more widely (*13,19*). The fact that we see relatively homogeneous spread of the presynthesized Xist RNPs during expansion, together with the observation that turnover occurs preferentially at the periphery of Xist RNA clusters, suggests individual Xist RNPs translocate and couple at successive anchor sites in a series of jumps (fig. S11 C). Further investigation of this aspect of Xist RNA behavior will require the development of new approaches that allow the movement of individual Xist RNPs to be monitored continually over time.

Our results demonstrate a key role for the silencing protein SPEN in long-range spreading of Xist RNA towards the chromosome termini. This novel function is independent of SPOC domain-mediated silencing activity, and hence provides an explanation for differences observed previously comparing SPEN null and SPOC domain mutant mESCs (*25*). Deletion of the Xist A-repeat, with which SPEN interacts, also results in disrupted spread of Xist RNA during expansion phase, as determined by RAP-seq analysis (*13*), and we suggest that this too is linked to the role of SPEN in Xist localization. An interesting possibility is that the SPOC-independent function of SPEN is important for transferring Xist RNA from initial 3D proximal sites to other locations on the chromosome. Although the underlying molecular mechanism for SPEN affecting Xist localization is unknown, our analysis of different mutations suggest a link to Xist RNP abundance and/or coupling. Which of these functions is important and whether or not they are interdependent remains to be determined.

## Acknowledgments

We thank Amanda Williams for assistance with NGS sequencing; Luke Lavis for kindly providing a range of HaloTag ligands; David Sherratt, University of Oxford, and Alfredo Castello, University of Oxford, for valuable discussions.

## Author contributions

N.B., L.S., H.C. and L.R. conceived experiments, L.R., H.C., J.B., B.R. and T.N. carried out experiments, N.B., L.S., L.R. and H.C. and wrote manuscript, L.R. and R.O. performed image processing and analysis, G.W., R.O. and L.R. ran computational analysis, R.O. and D.P. provided computational scripts.

## Data and code availability

All data, parameters and scripts used are available for reproducing the results of this investigation. Raw and reconstructed 3D-SIM images are available at the Image Data Resource (IDR). The programs required to run all scripts for ChaiN are: ImageJ (FIJI distribution with SIMcheck), R and Octave. Scripts for ChaiN are available in the following GitHub repositories: https://github.com/ezemiron/Chain. All the high throughput sequencing data encompassing ChrRNA-seq, ChIP-seq, and SLAM-seq are deposited in GEO under GSE154568.

## SUPPLEMENTARY MATERIALS

Materials and Methods

Figures S1-S11

Tables S1 and S2

