## supplemental text and figures for "Time-resolved structured illumination microscopy reveals key principles of Xist RNA spreading"

### SUPPLEMENTARY MATERIALS

### Material and Methods

#### *Generation of Cell lines*

*Tissue culture*

ESCs were grown in Dulbecco's modified Eagle's medium (DMEM; Life Technologies) supplemented with 10 % fetal calf serum (Seralab), 0.1 mM non-essential amino acids, 2 mM L-glutamine, 50 μM β-mercaptoethanol and 100 U/mL penicillin / 100 μg/mL streptomycin (Life Technologies), as well as 1000 U/mL LIF (made in-house). ESCs were grown on gelatin-coated plates with feeder cells, which were generated by inactivation of mouse fibroblasts with Mitomycin C (Sigma-Aldrich), at 37 °C with 5% $CO_2$ in a humid atmosphere. ESCs were passaged after treatment with TrypLE Express (Thermo Fisher Scientific) every 2-3 days. Before transfection, cells were grown on gelatin-coated 6-well plates with feeders.

*WT ES cell line*

iXist-Chr X ESCs were used as parental cell line for the generation of the WT ESC line (*26*). The BgI-stem-loop system was chosen for the labelling of Xist RNA. Due to concerns that BglG expression might be silenced over time, the BglG fusion protein was genetically engineered to be doxycycline inducible. Due to the superior fluorescent properties of organic dyes over fluorescent proteins and the possibility of flexible pulse labeling, a HaloTag approach was chosen for fluorescent labelling of the BglG protein for live cell and super-resolution imaging (*29*). Cells were transfected with 2 μg of a vector encoding for doxycycline inducible BglG-Halo fusion protein using Lipofectamine 2000 (Life Technologies) according to the manufacturer's instructions. Cells were then passaged to 90 mm gelatin-coated Petri dishes with feeders and grown in medium containing 200 μg/mL neomycin (Geneticin (G-418); Thermo Fisher Scientific) for 10 days. Clones were picked and genomic DNA (gDNA) was extracted, before successful insertions were screened by PCR (table S1). Selected clones were expanded and BglG-Halo expression was assessed by HaloTag ligand staining, and subsequent widefield fluorescence microscopy with an inverted fluorescence Axio Observer Z.1 microscope (Zeiss). Having selected a clone with intermediate BglG expression, low enough to minimize background but high enough to label all Xist molecules efficiently, a BgI-stem-loop array was inserted within exon 7 of the *Xist* locus by CRISPR/Cas9 mediated genome engineering. Here, a homology construct containing homology regions 1 kb 5′ and 3′ of the targeted insertion site within exon 7 of Xist was targeted by co-transfection with a specific sgRNA (table S1), which was designed using the WGE - CRISPR design tool (Wellcome Sanger Institute) and cloned into a backbone vector encoding for the CRISPR/Cas9 protein (PX459, Addgene). Cells were transfected overnight with 2 μg of the sgRNA construct and 1.15 μg of the targeting construct using Lipofectamine 2000 (Life Technologies) according to the manufacturer's instructions, and then passaged to 90 mm gelatinized Petri dishes with feeders. Cells were grown under antibiotic selection with 1.75 μg/mL puromycin for 48 h and thereafter grown for a further 9 d before being picked. Clones with successful insertion were selected by PCR screening after gDNA extraction (table S1). Successful labelling of BglG-Halo-Xist was confirmed by HaloTag ligand staining followed by 3D-SIM. Comparison of BglG-Halo-Xist signal intensity to background signal from unbound BglG-Halo allowed for the selection of the most suitable cell line for imaging experiments.

*Transgenic ES cell line*

XY P4D7 ESCs were used as parental cell line for the generation of the transgenic ESC line (*17*). A construct with doxycycline inducible Xist containing an array of 18 Bgl-stem-loops within exon 7 was used (*17*). Additionally, cells were transfected with the construct encoding doxycycline inducible BglG-Halo protein described earlier. Cells were co-transfected overnight with 5 µg of the transgenic Xist and 2.5 µg of the BglG-Halo construct using TransIT-LT1 transfection reagent (Mirusbio) according to the manufacturer's instructions. Cells were then passaged to 90 mm gelatin-coated Petri dishes with feeders and grown in medium containing 200 µg/mL neomycin (Geneticin (G-418); Thermo Fisher Scientific) for 10 d. Successful insertions of the BglG-Halo sequence were screened by PCR analysis of the gDNA (table S1). Transgenic Xist expression was assessed by Xist RNA FISH and subsequent widefield fluorescence microscopy with an inverted fluorescence Axio Observer Z.1 microscope (Zeiss). BglG-Halo expression and efficient labelling of transgenic BglG-Halo-Xist was confirmed by HaloTag ligand staining and 3D-SIM. Comparison of BglG-Halo-Xist signal intensity to background signal from unbound BglG-Halo allowed for the selection of the most suitable cell line for imaging experiments. Chromatin RNA sequencing later enabled determination of the insertion site of transgenic Xist on chromosome 15.

*Ciz-1 KO ES cell line*

The aforementioned WT ES cell line was used as parental cell line for the generation of the Ciz-1 KO ESCs by CRISPR/Cas9 mediated genome engineering. Here, a sgRNA, which had been previously used to induce a frameshift mutation (*10*), was used in combination with a second sgRNA designed using the WGE - CRISPR design tool to achieve a genomic deletion of 10.66 kb, resulting in the deletion of most of the *Ciz-1* gene (table S1). The sgRNAs were cloned into backbone vectors encoding for the CRISPR/Cas9 protein (PX459, Addgene). Cells were transfected overnight with 2 µg of each sgRNA construct using Lipofectamine 2000 (Life Technologies) according to the manufacturer's instructions. Cells were then passaged to 90 mm gelatinized Petri dishes with feeders and grown under antibiotic selection with 2 µg/mL puromycin for 48 h. Colonies were grown for 8 d before being picked. Selected clones with successful genomic deletion were screened for by PCR analysis of gDNA (table S1). Ciz-1 KO was confirmed by Western blot and immunofluorescence staining for Ciz-1, while successful labelling of BglG-Halo-Xist was confirmed by HaloTag ligand staining followed by 3D-SIM. A final clone for analysis was chosen based on the results of these assays, as well as the comparison of BglG-Halo-Xist signal intensity to background signal from unbound BglG-Halo.

*SPEN RRM del ES cell line*

The aforementioned WT ES cell line was used as parental cell line for the generation of the SPEN RRM del ESCs by CRISPR/Cas9 mediated genome engineering. Previously described sgRNAs (*21*) were used to achieve a genomic deletion of 38 kb, resulting in the KO of RRM2-4 of *Spen* (table S1). Cells were transfected and successful deletion was screened for as described above for the Ciz-1 KO cell line. SPEN RRM deletion in selected clones was confirmed by Southern blot. Presence of polyploid and XO cells was excluded by Xist RNA FISH and successful labelling of BglG-Halo-Xist was confirmed by HaloTag ligand staining followed by 3D-SIM. A final clone for analysis was chosen based on the results of these assays, as well as the comparison of BglG-Halo-Xist signal intensity to background signal from unbound BglG-Halo.

*SPOC domain mut ES cell line*

The aforementioned WT ES cell line was used as parental cell line for the generation of the SPEN SPOC mutant ESCs by CRISPR/Cas9 mediated genome engineering. The SPOC mutant, which disrupts the interaction between SPEN and NCoR/SMRT complex (*30*), was achieved by amino acid substitutions R3552A and R3554A in the SPOC domain of Spen. This substitution was previously described to abolish binding of Spen to NCOR1 an SMRT co-repressive complexes *in vitro* (*28*). The substitutions were engineered into a 0.5 kb homology construct corresponding to the region of Spen exon 14. The sgRNA (table S1) was cloned into a backbone vector encoding for the CRISPR/Cas9 protein (PX459, Addgene). Cells were transfected overnight with 1.42 $\mu$g of the sgRNA construct and 2.5 $\mu$g of the targeting construct using Lipofectamine 2000 (Life Technologies) according to the manufacturer's instructions. Cells were then passaged to 90 mm gelatinized Petri dishes with feeders and grown under antibiotic selection with 3 $\mu$g/mL puromycin for 48 h. Colonies were grown for 10 d before being picked. Selected clones with successful amino acid substitution were screened after gDNA extraction by NcoI digestion, PCR analysis and Sanger Sequencing of the PCR fragments (table S1). Presence of polyploid and XO cells was excluded by Xist RNA FISH and successful labelling of BglG-Halo-Xist was confirmed by HaloTag ligand staining followed by 3D-SIM. Chromatin RNA sequencing (ChrRNA-Seq) finally confirmed successful amino acid substitution and a final clone for analysis was chosen based on the cumulative results of these assays, as well as the comparison of BglG-Halo-Xist signal intensity to background signal from unbound BglG-Halo.

*NPC derivation*

In order to study Xist behavior in differentiated cells, WT and Ciz-1 KO ESCs were differentiated into neural progenitor cells (NPCs) following a following a previously adapted protocol (*31*,*32*). Cells were grown in gelatin-coated T25 flasks with feeder cells. Feeders were removed before differentiation by pre-plating three times for 40 min each. Then, 0.45x10$^6$ cells and 0.55x10$^6$ cells, respectively, were plated in a gelatin-coated T25 flasks each and grown in N2B27 medium containing 1 $\mu$g/mL doxycycline for 7 d. On day 7, the flasks were treated with Accutase (Millipore) and 3x10$^6$ cells each were plated in each 90 mm bacterial petri dishes with N2B27 medium containing 1 $\mu$g/mL doxycycline and 10 ng/mL EGF and FGF (Peprotech) to prevent cellular attachment. For imaging at day 10 of differentiation, cells were plated onto gelatin-coated 18x18 mm No. 1.5H precision coverslips (± 5 $\mu$m tol.; Marienfeld Superior) at day 7. At day 10, cell aggregates were collected by mild centrifugation, before being plated onto gelatin-coated 90 mm petri dishes in N2B27 medium containing 1 $\mu$g/mL doxycycline and 10 ng/mL EGF and FGF (Peprotech) each. When ~80 % confluent, cells were split 1:4 by Accutase (Millipore) treatment at room temperature followed by collection in PBS and centrifugation at 1,500 rpm for 5 min. For HaloTag staining at day 20 of differentiation, cells were plated onto precision coverslips at day 16 of differentiation. For Chromatin RNA Sequencing at day 30 of differentiation, cells were expanded onto one 145 mm petri dish per replica on day 25 of differentiation.

#### *Xist RNA FISH*

*Xist RNA FISH for scoring*

All cells were grown on gelatin-coated 18x18 mm No. 1.5H precision coverslips (± 5 μm tol.; Marienfeld Superior) in a 6-well plate on a layer of feeder cells. When reaching 60-70 % confluency, ESCs were induced for 3 h or 24 h with 1 μg/mL doxycycline, reserving one coverslip per experiment without doxycycline induction. After induction, coverslips were washed briefly with PBS twice, before being fixed with 3.7 % formaldehyde in PBS at room temperature for 10 min. Cells were rinsed with PBS and then permeabilized with 0.5 % Triton X-100 in PBS for 10 min at room temperature, followed by two washes with 70 % EtOH. Coverslips were then dehydrated with subsequent washes with 80 %, 95 %, 100 % EtOH for 5 min each, and dried briefly. Each coverslip was hybridized with 15 μL probe/ hybridization buffer mix. Xist probe was generated from an 18 kb cloned cDNA spanning the whole Xist transcript using a nick translation kit (Abbott Molecular) as previously described (*17*). 3 μL Texas Red labelled Xist RNA FISH probe per hybridization was added to 1/3 volume 10 mg/mL salmon sperm DNA, then precipitated by addition of 1/10 volume of 3 M NaOAc and 3 volumes 100 % EtOH. After washing with 70 % EtOH, the pellet was dried by speed vacuum and resuspended in 6 μL deionized formamide (Sigma) per hybridization. An equal amount of 2x hybridization buffer (4x SSC, 20 % dextran sulphate, 2 mg/mL BSA (NEB), 1/10 volume nuclease free water and 1/10 volume VRC pre-warmed at 65 °C for 5 min before use) was added, and the probe/ hybridization buffer mix was denatured at 75 °C for 5 min before being chilled on ice and used for hybridization in a humid chamber overnight at 37 °C. The next day, coverslips were washed 3 times with pre-warmed 50 % formamide/ 2x SSC at 42 °C for 5 min each, and subsequently 3 times with 2x SSC at 42 °C for 5 min each. Coverslips were mounted with 4′,6-diamidino-2-phenylindole (DAPI) containing Vectashield antifade mounting medium (Vector Labs) centrally on Superfrost Plus microscopy slides (VWR). Slides were dried, sealed using clear nail polish and cleaned for imaging. Imaging and scoring were carried out using an inverted fluorescence Axio Observer Z.1 microscope (Zeiss) using a PlanApo ×63/1.4 NA oil-immersion objective. Images were acquired using AxioVision software.

*Xist RNA FISH for 3D-SIM*

This protocol was adapted from the Stellaris FISH protocol (Biosearch Technologies) for a combination of Xist RNA FISH with HaloTag ligand staining of BglG-Halo-Xist. All cells were grown on gelatin-coated 18x18 mm No. 1.5H precision coverslips in a 6-well plate on a layer of feeder cells. When reaching 60-70 % confluency, ESCs were induced for 3 h or 24 h with 1 μg/mL doxycycline, before being stained with HaloTag ligand. After washout of the ligand, coverslips were washed twice with PBS and fixation was carried out by adding 3 % formaldehyde (pH 7) prepared in fresh PBS for 10 min at room temperature. A stepwise exchange of PBST (0.05 % Tween 20 PBS) was carried out, then coverslips were washed twice in PBST. Cells were permeabilized in 0.2 % Triton X-100 PBS for 10 min at room temperature, before being washed twice with PBST. Then, coverslips were blocked in 2 % BSA / 0.5 % fish skin gelatin / PBST containing freshly added RNAsin Plus (Promega) at a final concentration of 2 U/μL for 30 min. In the meantime, the Xist RNA FISH probe was prepared and precipitated as described before. However, after speed vacuum the probe pellet was resuspended in 12 μL Stellaris Hybridization buffer (Biosearch Technologies) per hybridization instead of formamide. The probe/hybridization buffer was prepared as described above, with the exception that the 2x hybridization buffer was prepared with 1/10 volume formamide before being added to the probe. After blocking, the cells were washed 3 times with PBST and fixed as before. Then, the coverslips were washed twice in PBST again, followed by a wash in 2x SSC. Next, the coverslips were incubated with FISH probe/ hybridization buffer overnight at 37 °C on parafilm in an extra humid chamber. The next day, the coverslips were transferred to a 6-well plate containing 1 mL wash buffer A (Biosearch Technologies) per well. After an incubation at 37 °C for 30 min, wash buffer A was replaced with 1 mL wash buffer A containing 5 ng/mL DAPI, and coverslips were incubated at 37 °C for 30 min. 1 mL of 2x SSC was added to each well, followed by an incubation of 5 min at room temperature. Finally, coverslips were mounted centrally on the unfrosted side of Superfrost Plus microscopy slides (VWR) using Vectashield antifade mounting media (w/o DAPI), sealed with clear nail polish and imaged within a week using the DeltaVision OMX V3 Blaze system (GE Healthcare).

***Immunofluorescence labelling***

All cells were grown on gelatin-coated 18x18 mm No. 1.5H precision coverslips in a 6-well plate on feeder cells. When reaching 60-70 % confluency, ESCs were induced for 3 h or 24 h with 1 μg/mL doxycycline. Optionally, cells were stained with HaloTag ligand at this stage for a combination of HaloTag ligand and immunofluorescence staining. Coverslips were washed twice with PBS and fixation was carried out by adding 2 % formaldehyde prepared in fresh PBS (pH 7) for 10 min at room temperature. A stepwise exchange of PBST was carried out, before coverslips were washed twice in PBS. Then, cells were permeabilized with 0.2 % Triton X-100 in PBS for 10 min at room temperature, followed by a wash with PBST. Coverslips were blocked using 3 % BSA/ 5 % normal goat serum/ PBST on parafilm in a humid chamber for 30 min at room temperature, and then incubated with primary antibody (table S2) diluted in block buffer for 1 h in a humidified chamber at 37 °C. After 3 washes in PBST, coverslips were incubated with the Alexa Fluor-conjugated secondary antibody (table S2) diluted 1:1000 in block buffer for 30 min in a humidified chamber at 37 °C. After another 4 washes with PBST, the cells were post-fixed as before, followed by a final wash in PBST. Lastly, coverslips were incubated with 2 μg/mL DAPI in PBS for 10 min, before being mounted in Vectashield centrally on the unfrosted side of Superfrost Plus microscopy slides (VWR) using Vectashield soft mount media, sealed with clear nail polish and imaged within a week using the DeltaVision OMX V3 Blaze system (GE Healthcare).

***HaloTag staining***

All cells were grown on gelatin-coated 18x18 mm No. 1.5H precision coverslips in a 6-well plate on feeder cells. When reaching 60-70 % confluency, ESCs were induced for 3 h or 24 h with 1 μg/mL doxycycline. Then, cells were incubated with 50 nM diAcFAM HaloTag ligand (488 nm, Promega) and 1 μg/mL doxycycline for 45 min, before being washed with medium containing 1 μg/mL doxycycline for 15 min. After a brief wash with PBS, cells were fixed for 10 min at room temperature with 2 % formaldehyde prepared fresh in PBS (pH 7). A stepwise exchange of PBST was carried out, and cells were permeabilized with 0.2 % Triton X-100 for 10 min, followed by two washes with PBST. Next, cells were incubated with 2 $\mu$g/mL DAPI in PBST for 10 min, before being washed briefly with PBS and, subsequently with dd$H_2O$ to remove salt, and equilibrated with Vectashield mounting medium. The coverslips were mounted centrally on the unfrosted side of Superfrost Plus microscopy slides (VWR) using Vectashield soft mount media, sealed with clear nail polish and imaged within a week using the DeltaVision OMX V3 Blaze system (GE Healthcare).

***3D-SIM***

*Acquisition*

3D-SIM imaging was performed on a DeltaVision OMX V3 Blaze system (GE Healthcare) equipped with a 60x/1.42 NA Plan Apo oil immersion objective (Olympus), pco.edge 5.5 sCMOS cameras (PCO), and 405, 488, 593 and 640 nm lasers. Image stacks were acquired with a z-distance of 125 nm and with 15 raw images per plane (5 phases, 3 angles). Spherical aberration after reconstruction was reduced by using immersion oil of different refractive indices (RIs) matched to respective optical transfer functions (OTFs). Here, immersion oil with an RI of 1.514 was used for the sample acquisition and matched to OTFs generated using immersion oil of RI 1.512 for the blue and green, and 1.514 for the red channel. OTFs were acquired using 170 nm diameter blue emitting PS-Speck beads and 100 nm diameter green and red emitting FluoSphere beads (Thermo Fisher Scientific).

*Reconstruction*

The raw data was computationally reconstructed with softWoRx 6.5.2 (GE Healthcare) using channel-specific OTFs and Wiener filter settings of 0.005. A lateral (x-y) resolution of approximately 120 nm and an axial (z) resolution of approximately 320 nm was achieved (*33*). All data underwent assessment via SIMcheck (*34*) to determine image quality via analysis of modulation contrast to noise ratio (MCNR), spherical aberration mismatch, reconstructed Fourier plot and reconstructed intensity histogram values. Of note, DAPI reconstructions in mESC were typically below the quality threshold levels of average MCNR of 5, and were therefore only used to indicate the nuclear outlines. Reconstructed 32-bit 3D-SIM datasets were thresholded to the stack modal intensity value and converted to 16-bit composite z-stacks to discard negative intensity values using SIMcheck's "threshold and 16-bit conversion" utility and MCNR maps were generated using the "raw data modulation contrast" tool of SIMcheck. To eliminate false positive signals from reconstructed noise, we applied an MCNR filtering using an adapted in-house ImageJ script (*33*). Here, all pixels in the reconstructed dataset where the corresponding MCNR values in the raw data map fall below an empirically chosen threshold of 4 are set to zero. Thereafter, the resulting 'masked' reconstructed dataset is blurred with a Gaussian filter with 0.8 pixel radius to smoothen hard edges (fig. S2B).

*Alignment*

Color channels were registered in 3D with the open-source software Chromagnon 0.85 (*35*) determining alignment parameter (x,y,z-translation, x,y,z-magnification, and z-rotation) from a 3D-SIM datasets acquired on the date of image acquisition of multicolor-detected EdU pulse replication labelled C127 mouse cells serving as biological 3D alignment calibration sample (*20*).

*3D-SIM live cell imaging*
Cells were grown on gelatin-coated 35 mm No. 1.5H glass bottom imaging dishes (Ibidi) on feeder cells. When reaching 50 % confluency, ESCs were induced for 24 h with 1 μg/mL doxycycline. Then, cells were incubated with medium containing 50 nM JF-646 HaloTag ligand (Janelia Fluor 646, kindly provided by Luke Lavis, HHMI Janelia) and 1 μg/mL doxycycline for 45 min, before being washed with medium containing 1 μg/mL doxycycline for 15 min. Before imaging, cells were washed briefly with PBS and medium was replaced with phenol red free medium with 1 μg/mL doxycycline. For live-cell 3D-SIM image stacks were acquired in 10 s intervals for up to 10 min. Image acquisition was performed at 37 °C with 5 % $CO_2$. To adapt for the temperature increase, immersion oil with an RI of 1.520 was used for the sample acquisition and matched to OTFs generated. Images were then processed as described above, with the exception that bleach correction was carried out.

***RNA-SPLIT***
*Sample preparation*
All cells were grown on gelatin-coated 18x18 mm No 1.5H precision coverslips as described above for HaloTag staining. ESCs were induced for different times with 1 μg/mL doxycycline and stained with HaloTag ligands dependent on the variation of RNA-SPLIT as described below. RNA-SPLIT was determined to achieve optimal results when using the diAcFAM HaloTag ligand (Promega) as first pulse label and the JF-585 HaloTag ligand (kindly provided by Luke Lavis, HHMI Janelia) as second pulse label. Initial experiments with different dye concentrations showed that a concentration of 50 nM for each of the ligands yielded optimal results. After a wash with PBS, samples were fixed and mounted as described above.

*Nascent Xist RNA dynamics*
For assessing the transcription dynamics of Xist RNA, cells grown on precision coverslips were sequentially stained with two different HaloTag ligands in each experiment. For analysis of expansion or steady state phase, cells were induced for either 1.5 h or 24 h with 1 μg/mL doxycycline, respectively. Then, cells were incubated with 50 nM diAcFAM HaloTag ligand (488 nm, Promega) and 1 μg/mL doxycycline for 45 min to label pre-synthesized Xist RNA molecules, before washing with medium containing 1 μg/mL doxycycline for 15 min. Next, the different coverslips were incubated with 50 nM JF-585 HaloTag ligand and 1 μg/mL doxycycline for 10 min, 20 min, 30 min, 40 min, 50 min and 60 min respectively to label newly synthesized Xist RNA molecules, before being washed with PBS. Induction and staining times with the first HaloTag ligand were staggered, such that all coverslips could be fixed at the same time despite the different staining times with the second HaloTag ligand.

*Xist turnover on chromatin*
For assessing the turnover of Xist RNA, cells grown on precision coverslips were stained with 50 nM diAcFAM HaloTag ligand after 1.5 h or 24 h doxycycline induction, respectively, as described above. After the first pulse, coverslips were incubated with 50 nM JF-585 HaloTag

ligand and 1 μg/mL doxycycline for 60 min, 80 min, 100 min, 120 min, 140 min, 160 min, 180 min, 200 min and 220 min respectively to label newly synthesized Xist RNA molecules, while one coverslip per experiment was not stained with the second HaloTag ligand at all and served as 0 min time point. Coverslips were finally washed with PBS. Induction and staining times with the first HaloTag ligand were staggered, such that all coverslips could be fixed at the same time despite the different staining times with the second HaloTag ligand. For each experiment, the endpoint was determined as when signal from the first HaloTag ligand could no longer be detected in the cells for two timepoints in a row.

*Image analysis workflow*

After image acquisition, reconstruction and pre-processing as described earlier, images were further processed and subsequently analyzed. First, signal with low modulation contrast (MCN) was discarded by an adapted in-house ImageJ script based on the MCNR maps and thresholded files generated, as described above. The MCN threshold was set at 4 with a Gaussian blur of 0.8 pixel radius. Next, manual thresholding of the red and green channel was performed in order to discard background signal originating from free diffusing fluorescent BglG-Halo. Thresholding was performed conservatively to limit detection to true- positive signal. Lateral color channel alignment was performed as described above. The resulting images were used as representative images of whole nuclei. For further analysis however, the DAPI channel was discarded, and Xist territories were cropped manually using ImageJ to exclude signal from BglG-Halo accumulation in the nucleoli, or signal from other cells. The cropped dimensions were later used to define Xist territory volume during expansion and steady state in all different cell types.

The resulting image files were subsequently analyzed using an in-house adapted *makefile* script for masking of the signal and centroid determination by watershed algorithm, which also allowed for the determination and comparison of the intensities of different Xist foci. The output data was used to determine the number of Xist foci during expansion and steady state phase in all different cell types. Moreover, it was used to determine the transcription rates based on the centroid count of the HaloTag ligand labelling newly synthesized Xist RNA molecules, as well as the turnover of Xist RNA based on the count of the HaloTag ligand labelling pre-synthesized Xist RNA molecules.

Nearest neighbor analysis (NNA) and variations thereof were conducted based on the x, y and z coordinates of centroids determined previously using a different in-house adapted *makefile* script. Distances determined from HaloTag ligand labelling newly synthesized RNA to the HaloTag ligand labelling pre-synthesized RNA were used to assess pairing behavior of Xist RNA during expansion and steady state in all cell types. Data from female C127 cells pulsed with EdU used previously for alignment was used as a technical control to determine nearest neighbor distances of molecules labelled with different ligands at the same time. Here, only EdU files acquired on the days of corresponding RNA-SPLIT data acquisition were used, allowing for sample-specific matched technical controls. Additionally, 'Random' sample controls were generated in a sample-specific manner by scrambling of the x, y and z coordinates of the centroids determined for newly synthesized RNA of each specific sample using Microsoft Excel.

Using a variation of the aforementioned NNA, median distances between each molecule and its ten nearest neighboring molecules in the same channel were determined as a measure of Xist RNA density. Combined densities of newly synthesized and pre-synthesized RNA in expansion and steady state were used for comparison of Xist RNA densities in different cell types. For NPCs in particular, median distances between each molecule and all other molecules were determined as measure of Xist RNA density in order to account for clustering of Xist RNA at different sites throughout the nucleus in Ciz-1 KO NPCs.

Approximate localizations of the Xist transcription site were determined for each Xist territory based on fluorescence intensity and density of the newly synthesized Xist RNA signal of WT ESCs. A variation of NNA was used to measure the distance of said approximate transcription site to all newly synthesized Xist molecules and to all pre-synthesized Xist molecules separately. In order to account for variability in Xist territory size, distances were normalized to the maximum distance measured for each territory, which was set to 1. This was then used to characterize the spreading behavior of 'new' Xist RNA molecules by comparing distance distributions in relation to the transcription site over time. Here, the Xist territory was divided into 3 zones, with zone 1, 2 and 3 extending 0-50 %, 50-75 % and 75-100 % of the maximum distance from the transcription site respectively. Then, the proportion of newly synthesized or pre-synthesized Xist RNA molecules in each zone over time was quantified. This approach allowed for the time-resolved comparison of newly synthesized Xist RNA molecule and pre-synthesized Xist RNA molecule distributions during both expansion and steady state in WT ESCs. A rate of expansion could be calculated, since NNA distances were measured in 3D and the 3D volumes of the clusters were already known. On average, Xist clusters had a size of 98.9 $\mu m^3$ in expansion, which can be set equal to 100 % of the normalized volume used to determine Xist spreading previously. On average, newly synthesized Xist extends across 29.9 % of this volume after 20 min and 35.2 % after 60 min, meaning that there is an increase of 5.3 % of the Xist expansion volume within 40 min. 5.3 % of the average Xist cluster volume of 98.9 $\mu m^3$ corresponds to 5.11 $\mu m^3$. Based on this, the expansion rate of newly synthesized Xist RNA can be approximated as 7.7 $\mu m^3$ $h^{-1}$. Furthermore, the localizations of transcription sites could be used to assess their distribution during expansion as compared to steady state phase. Here, 3D coordinates of Xist transcription sites were normalized to Xist territory dimensions and Xist territories were divided into 3 zones to account for variability in Xist territory size.

*Dual-color control*

In order to exclude that the results obtained by NNA are result from simultaneous labelling of Xist RNA molecules with BglG-Halo fusion proteins labelled with the two different HaloTag ligands, a dual color control was performed. A total of 6 coverslips with WT ESCs were induced for 24 h with 1 $\mu$g/mL doxycycline, before being incubated with medium containing 50 nM diAcFAM HaloTag ligand (488 nm, Promega), 50 nM JF-585 HaloTag ligand and 1 $\mu$g/mL doxycycline for 45 min. Then, coverslips were washed with medium containing 1 $\mu$g/mL doxycycline for 15 min, followed by a wash with PBS and preparation for imaging as described previously.

*Dox washout experiment*

In order to assess the turnover on chromatin of Xist RNA when Xist transcription is reduced, a washout of doxycycline was performed. Here, 10 coverslips with cells were sequentially stained with two different HaloTag ligands. WT ESCs were induced for 1.5 h with 1 $\mu$g/mL doxycycline.

Cells were then incubated with medium containing 50 nM diAcFAM HaloTag ligand (488 nm, Promega) but without doxycycline for 45 min to label pre-synthesized Xist RNA molecules, before washing just with medium for 15 min. Next, the different coverslips were incubated with 50 nM JF-585 HaloTag ligand but without doxycycline for 60 min, 80 min, 100 min, 120 min, 140 min, 160 min, 180 min, 200 min and 220 min respectively to label newly synthesized Xist RNA molecules, while one coverslip was not stained with the second HaloTag ligand and served as the 0 min time point. Coverslips were washed with PBS and samples were prepared for imaging as described previously. Induction and staining times with the first HaloTag ligand were staggered, such that all coverslips could be fixed at the same time despite the different staining times with the second HaloTag ligand. Successful reduction of Xist transcription was determined by comparison of the newly synthesized Xist RNA molecule count determined for the doxycycline washout experiment to that determined for WT ESCs during expansion.

***SLAM-Seq***

*Sample preparation*

SLAM-seq of WT cells, as well as the corresponding parental cell line with untagged Xist, was performed using the SLAMSeq Kinetics Kit, Catabolic Kinetics Module (Cat. No. 062.24, Lexogen). Cells were grown in gelatin-coated 6-well plates after pre-plating to discard feeder cells, with 6 wells needed for each experimental replica. When reaching 60-70 % confluency, cells were induced with 1 $\mu$g/mL doxycycline for 17.5 – 20 h depending on the length of the subsequent washout times. Then, transcribed RNA was labelled with 4SU by incubation with medium containing 500 $\mu$M 4SU (Lexogen) and 1 $\mu$g/mL doxycycline for 4 h. Next, 4SU was withdrawn by washout for all samples except the 0 min sample. Different samples were washed with medium containing 1 $\mu$g/mL doxycycline and 50 mM uridine (in excess, Lexogen) for 30 min, 60 min, 90 min, 120 min and 150 min respectively. Induction, 4SU labelling and withdrawal times were staggered, such that all samples could be harvested at the same time despite the different washout times. After cells were collected by centrifugation at 1,500 g for 5 min in PBS, nuclear extracts were obtained as follows: Cells were washed again with PBS and resuspended in 10 volumes buffer A (10 mM HEPES pH 7.9, 1.5 mM $MgCl_2$, 10 mM KCl, 0.5 mM DTT and complete protease inhibitors in RNase free water). Subsequently cells were incubated on ice for 10 min and recovered by centrifugation, followed by resuspension in 3 volumes buffer A with 0.1 % NP-40 for 10 min. Nuclei were recovered by centrifugation at and TRIzol/chloroform RNA extraction was performed using the SLAMSeq Kinetics Kit (Lexogen). Then, samples were treated with Iodoacetamide to modify the 4-thiol group of S4U-containing nucleotides via the addition of a carboxyamidomethyl group using the SLAMSeq Kinetics Kit (Lexogen). The RNA was precipitated and washed again, before being resuspended in nuclease free water. 1 ng RNA of each sample were run on the bioanalyzer to determine the exact RNA concentration and RNA integrity of all samples. Based on this, 1 $\mu$g RNA of each sample was then taken forward for library preparation using the Illumina TruSeq stranded total RNA kit (RS-122-2301). Quantification of the libraries was performed by qPCR using KAPA Library Quantification DNA standards (Kapa Biosystems, KK4903). Finally, the libraries were pooled and 2×81 paired-end sequencing was performed using Illumina NextSeq500 (FC-404-2002).

*Data analysis*
Estimation of nuclear RNA half-life was performed using GRAND-SLAM (*36*). Briefly, the paired-end sequencing reads were mapped to mouse genome mm10 by STAR (v2.5.2b) (*37*) with the key parameters (--outFilterMultimapNmax 1 --outFilterMismatchNmax 999 --alignEndsType EndToEnd). Given that T to C conversion is the signature of SLAMSeq, the T to C conversion rate was calculated as 4SU incorporation and the average of all the rest of the conversions was calculated as background due to errors from sequencing or library preparation. The T to C conversion rate and background rate were calculated accordingly for each sample. The nuclear RNA decay was assumed to follow an exponential model, so the T2C conversion rate (or T2C conversion rate-corresponding background rate) was fitted to the exponential model to estimate the RNA's nuclear half-life. In addition, the sample (dox-treated A11B2 cells, ChrRNA-Seq) (*26*) without 4SU incorporation was used to estimate the background mutation rate for all 12 potential mismatches. Half-life of Xist RNA and another nuclear lncRNA Pvt1 were plotted.

***Chromatin RNA Sequencing***
*Sample preparation*
For chromatin RNA sequencing experiments, cells were grown in gelatin-coated T25 flasks with feeder cells. When confluent, cells were pre-plated for 40 min twice to discard the feeder cells, and expanded to gelatin-coated 15 cm dishes. Before chromatin RNA extraction, cells were induced for 3 h or 24 h with 1 $\mu$g/mL doxycycline for expansion and steady state respectively. A control sample comprising cells without addition of doxycycline was included with the analysis of each cell line. Chromatin RNA for each sequencing replica was extracted from one confluent 15 cm dish of cells. Cells were harvested and recovered by centrifugation at 1,000 rpm for 3 min at room temperature, before being snap frozen on dry ice. Next, pellets were resuspended in RLB (10 mM Tris pH 7.5, 10 mM KCl, 1.5 mM $MgCl_2$, and 0.1 % NP40) and incubated on ice for 5 min to lyse the cells. Nuclei were purified by centrifugation through a sucrose cushion (24 % sucrose in RLB) at 2,800 g for 10 min at 4 °C. The pellets were resuspended in NUN1 (20 mM Tris pH 7.5, 75 mM NaCl, 0.5 mM EDTA, 50 % glycerol), and subsequently lysed with NUN2 (20 mM HEPES pH 7.9, 300 mM NaCl, 7.5 mM $MgCl_2$, 0.2 mM EDTA, 1 M urea). Samples were incubated for 15 min on ice, vortexing occasionally, before being centrifuged at 2,800 g for 10 min at 4 °C to isolate the insoluble chromatin fraction. The pellets were resuspended in TRIzol and homogenized by passing multiple times through a 23 gauge needle. Finally, chromatin-associated RNA was purified through standard TRIzol/chloroform extraction followed by isopropanol precipitation. Samples were treated with Turbo DNase and the RNA was purified by RNeasy column cleanup (Qiagen). 100 ng - 1 $\mu$g RNA of each sample were taken forward for library preparation using the Illumina TruSeq stranded total RNA kit (RS-122-2301). Quantification of the libraries was performed by qPCR with KAPA Library Quantification DNA standards (Kapa Biosystems, KK4903). Finally, the libraries were pooled and 2 × 81 paired-end sequencing was performed using Illumina NextSeq500 (FC-404-2002).

*Data analysis*
The analysis strategy that was used for chromatin RNA sequencing has been described in detail (*26*). In brief, raw fastq files of read pairs were mapped to an rRNA build by bowtie2 (v2.3.2), while rRNA-mapped reads were discarded. Remaining unmapped reads were aligned to the "N-

masked” genome (from mm10 coordinates) with STAR (v2.4.2a) using parameters “–outFilterMultimapNmax 1 –outFilterMismatchNmax 4 –alignEndsType EndToEnd”. Unique alignments were retained for further analysis. 23,005,850 SNPs between Cast and 129S genomes identified previously (38), were used for allelic split. These were used to split the alignment into distinct alleles (Cast and 129S) with the help of SNPsplit (v0.2.0; Babraham Institute, Cambridge, UK). The allelic read numbers were counted using the program featureCounts (-t transcript -g gene_id -s 2) (39), while the alignments were sorted by Samtools (40). For bi-allelic analysis, counts were normalized to one million mapped read pairs using the edgeR R package. Genes with a minimum of 10 SNP-covering reads across all the samples were further taken to calculate the allelic ratio of Xi/(Xi+Xa). Here, Xi and Xa indicate the inactive and active allele, respectively. Xist mediated gene silencing during expansion and steady state was determined by the difference in the aforementioned allelic ratios between uninduced and doxycycline induced samples.

#### *ChIP-seq*

*ES Tissue Culture*

Biological replicate clones for SPEN RRM del and SPOC domain mutants were derived from iXist-Chr X cells. ES cells cultured on feeders were pre-plated for 30 min, then plated on 15 cm dishes. Xist was induced by addition of 1 $\mu$g/mL doxycycline to growth media 3 h or 24 h prior to harvesting cells. Whereas all samples for the 3 h experiment were processed in parallel on the same day, for the 24 h experiment the SPEN RRM del lines, SPOC domain mut lines, and each of the three replicates of iXist-Chr X (WT) cells were processed on separate occasions.

*Native ChIP-seq*

H2AK119ub1 native ChIP-seq was performed for iXist-Chr X, SpenRRM and SpocMut cell lines largely as previously described (26) using buffers supplemented with 10 mM of the deubiquitinase inhibitor N-ethylmaleimide (Sigma, E3876-5G) throughout. Briefly, $50x10^6$ ES cells were lysed in RSB (10 mM Tris pH 8, 10 mM NaCl, 3 mM $MgCl_2$, 0.1 % NP40) for 5 min on ice with gentle inversion. Nuclei were resuspended in 1 mL of RSB + 0.25 M sucrose + 3 mM $CaCl_2$, treated with 200 U of MNase (Fermentas) for 5 min at 37 °C, quenched with 4 $\mu$L of 1 M EDTA, then centrifuged at 5,000 rpm for 5 min. The supernatant was transferred to a fresh tube as fraction S1. The chromatin pellet was incubated for 1 h in 300 $\mu$L of nucleosome release buffer (10 mM Tris pH 7.5, 10 mM NaCl, 0.2 mM EDTA), carefully passed five times through a 27G needle and then centrifuged at 5 000 rpm for 5 min. The supernatant from this S2 fraction was combined with S1 as soluble chromatin extract. For each ChIP reaction, 100 $\mu$L of chromatin was diluted in Native ChIP incubation buffer (10 mM Tris pH 7.5, 70 mM NaCl, 2 mM $MgCl_2$, 2 mM EDTA, 0.1 % Triton) to 1 mL and incubated with 4 $\mu$L H2AK119ub1 Ab (Cell Signalling Technology, #8240) overnight at 4 °C. Samples were incubated for 1 h with 40 $\mu$L protein A agarose beads pre-blocked in Native ChIP incubation buffer with 1 mg/mL BSA and 1 mg/mL yeast tRNA, then washed for a total of four times in Native ChIP wash buffer (20 mM Tris pH 7.5, 2 mM EDTA, 125 mM NaCl, 0.1 % Triton) and once in TE. The DNA was eluted with 1 % SDS and 100 mM $NaHCO_3$, and was purified using the ChIP DNA Clean and Concentrator kit (Zymo Research). 50 ng of ChIP DNA was used for library prep using the NEBNext Ultra II DNA Library Prep Kit with NEBNext Single indices (E7645), and then quantified using a

Bionalayzer 2100 (Agilent) and a Qubit fluorometer (Invitrogen). The libraries were pooled and 2×81 paired-end sequencing was performed using Illumina NextSeq500 (FC-404-2002).

*Native ChIP-seq data analysis*

Raw fastq read pairs were mapped to the "N-masked" mm10 genome by bowtie2 (v2.2.6) using parameters "--very-sensitive --no-discordant --no-mixed -X 2000". Alignment files were subsequently filtered to remove unmapped read pairs and PCR duplicates (picard-tools MarkDuplicates). For allelic analysis, we made use of SNPs between Cast and 129S genomes and employed SNPsplit (v0.2.0; Babraham Institute, Cambridge, UK) to split 55-60 % of filtered alignments into distinct alleles (Cast and 129S) using the parameter "--paired". Each alignment file was then processed into bedGraph format by genomeCoverageBed (BEDtools, *41*) and normalized to the total library size of the sample. The custom script ExtractInfoFrombedGraph_AtBed.py (https://github.com/guifengwei) was used to extract values of H2AK119ub1 enrichment for 250 kb windows spanning the whole X chromosome (Fig. 5) or the 103.5 Mb region of Chr X proximal to Xist (fig. S11). Gain of H2AK119ub1 upon Xist expression was calculated by subtraction of Uninduced from 24 h or 3 h induced samples (Dox – NoDox). Line plots were generated in R from replicates averaged for each mutant (3x iXist-Chr X technical replicates, 2x Spen KO clones, 2x SPOC mutant clones).

***Western blot***

For Ciz-1 Western blot experiments, the candidate KO clones, and WT cells serving as control, were grown in gelatin-coated T25 flasks each with feeder cells. When confluent, cells were pre-plated for 40 min twice to discard the feeder cells, and subsequently plated in a well of a gelatin-coated 6-well plate each. When confluent, cells were washed with PBS and detached using TrypLE Express (Thermo Fisher Scientific). Cells were washed and recovered by centrifugation, before $3x10^6$ cells of each clone were dissolved in 200 $\mu$L 2x SMASH buffer (33 mM Tris-HCl (pH 6.8), 11 % Glycerol, 40 mg/mL SDS, 200 $\mu$g/mL bromophenol blue and 10 % β-mercaptoethanol in nuclease free water). Samples were left shaking for 20 min, and then snap frozen on dry ice and thawed once. Samples were then separated by SDS polyacrylamide gel electrophoresis (8 % separating gel and 5 % stacking gel) at 80 V for 15 min, and then at 180V for another 70 min. Next, samples were transferred onto a PVDF membrane by semi-dry transfer at 15 V for 1 h. The membrane was blocked by incubation with 10 mL PBS, 0.1 % Tween (PBST) with 5 % w/v Marvel milk powder for 1 h at room temperature. The membrane was then incubated with the corresponding primary antibody (Ciz-1, anti-rabbit, affinity purified, polyclonal (a kind gift from the Coverley lab, University of York); and tubulin, anti-rabbit, Cell Signaling, Cat. No 2144) diluted 1:1000 in PBST with 5 % w/v Marvel milk powder at 4 °C overnight. The membrane was washed 3 times in PBST with 5 % w/v Marvel milk powder for 10 min each, before being incubated with secondary antibody conjugated to horseradish peroxidase (anti-rabbit, HRP linked, Amersham) diluted 1:1000 in PBST with 5 % w/v Marvel milk powder for 1 h. After washes with PBST with 5 % w/v Marvel milk powder and finally with PBST followed by PBS only, bands were visualized using ECL (GE Healthcare) for antibody detection.

#### *Southern blot*

For Southern blot analysis of candidate Spen RRM KO clones, cells were grown in gelatin-coated T25 flasks each with feeder cells. When confluent, cells were pre-plated for 40 min twice to discard the feeder cells, and subsequently plated in a well of a gelatin-coated 6-well plate each. Cells were washed 3 times with PBS, before 1 mL lysis buffer (10 mM NaCl, 10 mM Tris pH 7.5, 10 mM EDTA pH 8, 0.5 % sarcosyl; filter sterilized; 1 mg/mL Proteinase K added fresh) was added directly to each well to lyse the cells completely. Lysates were incubated at 55 °C shaking at 300 rpm overnight. Standard phenol chloroform extraction of the DNA was carried out using MaXtract High Density tubes (Qiagen). DNA was precipitated with NaCl and EtOH and the DNA was recovered by were centrifugation. Next, an EcoRV restriction digest of the samples was performed and 5 $\mu$g digested DNA each were run on a 1 % EtBr agarose gel for ~4 h at 50 V. The gel was incubated with depurination solution (0.125 M HCl in MiliQ water) and for 15 min, with denaturation buffer (0.5 M NaOH, 1.5 M NaCl in Milli-Q water) for 45 min and finally with neutralization buffer (0.5 M Tris, 1.5 M NaCl in Milli-Q water, pH 8) for 30 min. After capillary transfer overnight to an Immobilon NY+ membrane (Millipore) for at least 18 h, the membrane was air dried and UV crosslinked using the Stratalinker.

The membrane pre-hybridized with Dextran - SLS - SCC hybridization solution pre-warmed to 65 °C (83 mg/mL dextran sulphate, 5x SSC and 0.875 % N-lauryl sarcosine sodium salt solution in MilliQ water filtered through a 0.8 $\mu$M membrane before addition of 48 $\mu$g/mL denatured sheared salmon sperm DNA just before use) for 1 h at 65 °C. The probe was prepared using the prime-H II kit (Stratagene) as follows from a 626bp fragment downstream of the sgRNA used for the Spen RRM KO in exon 12 (table S1). The probe was purified using a G50 Micro column (GE Healthcare), before being boiled for 5 min at 95 °C. The denatured probe was placed on ice for 1 min, added to the hybridization mixture and membrane, and then hybridized overnight at 65 – 67 °C. The next day, the membrane was washed twice with low stringency at 65 °C for 15 min each. It was subsequently exposed in a Fuji cassette overnight, before it was scanned on a FLA-7000 (Fujifilm). Mutant bands were detectable at 2.64 kb, while WT bands were detectable at 4 kb.

#### *Statistical analysis*

For the comparison of two independent large datasets throughout the project, statistical significance was determined by conducting an unpaired two-sample Wilcoxon test using R. This statistical test determines whether it is equally likely that a randomly selected value from one dataset will be less than or greater than a randomly chosen value from another population, making it ideal to determine whether two large datasets have the same distribution. Hence, the unpaired two-sample Wilcoxon test was used as a non-parametric alternative to the unpaired t-test to determine statistical significance.

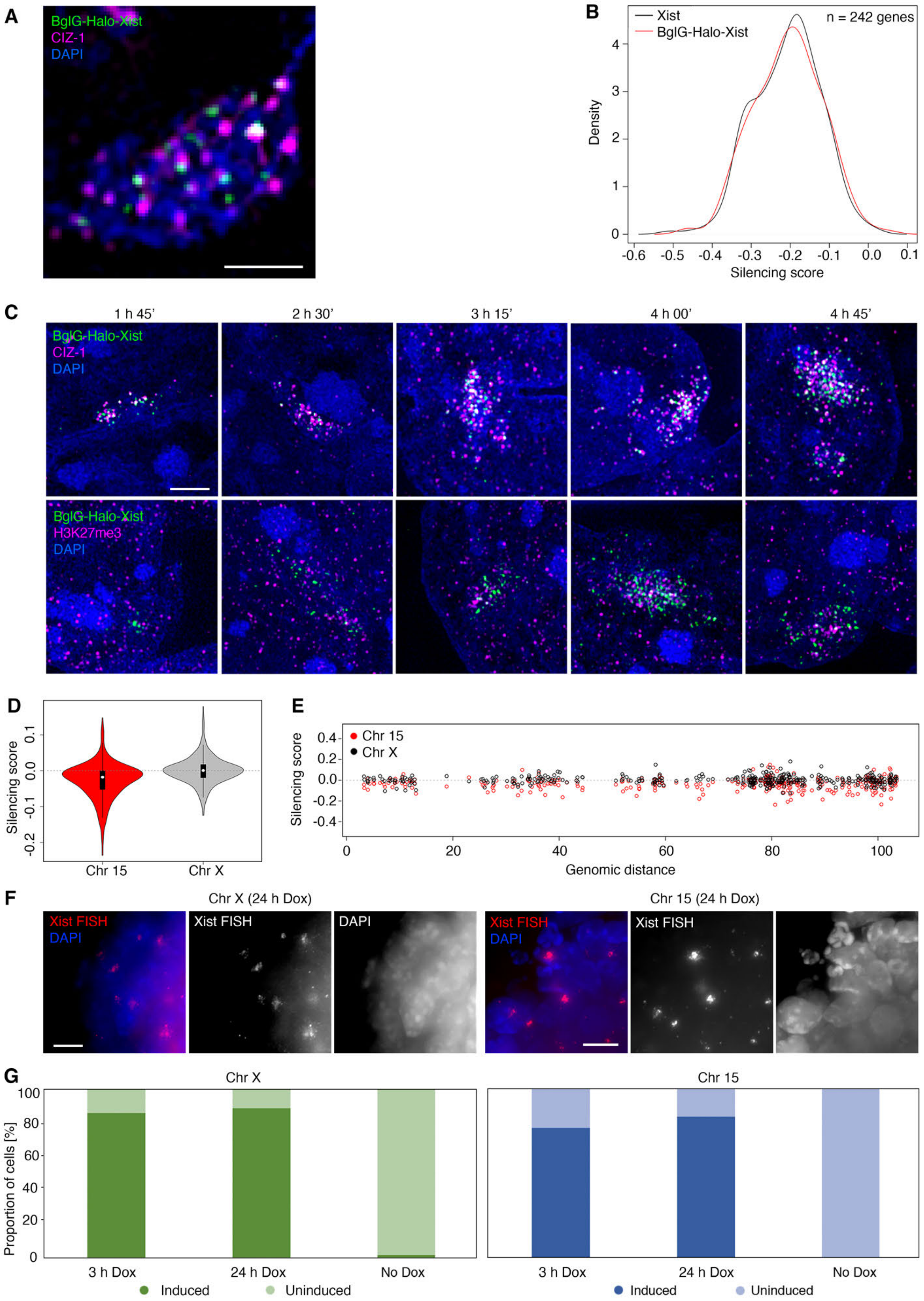
A
BglG-Halo-Xist
CIZ-1
DAPI
B
Xist
BglG-Halo-Xist
n = 242 genes
Density
Silencing score
C
1 h 45'
2 h 30'
3 h 15'
4 h 00'
4 h 45'
BglG-Halo-Xist
CIZ-1
DAPI
BglG-Halo-Xist
H3K27me3
DAPI
D
Silencing score
Chr 15
Chr X
E
Chr 15
Chr X
Silencing score
Genomic distance
F
Chr X (24 h Dox)
Xist FISH
DAPI
Xist FISH
DAPI
Chr 15 (24 h Dox)
Xist FISH
DAPI
Xist FISH
G
Chr X
Chr 15
Proportion of cells [%]
3 h Dox
24 h Dox
No Dox
Induced
Uninduced

**Fig. S1. Characterization of BglG-Halo-Xist functionality on Chr X and Chr 15.**
(**A**) Representative 3D-SIM image (single z-section) showing immunostaining of Ciz-1 in combination with Xist RNA HaloTag staining and DAPI counterstaining. Scale bar: 1 $\mu$m. (**B**) Plot representing X-linked gene silencing induced by BglG-Halo tagged (red) and untagged (black) Xist RNA after 24 h of doxycycline induction. (**C**) Representative 3D-SIM images (z-projections) illustrating Halo-Tag labelled Xist RNA with immunostaining of Ciz-1 (top) and H3K27me3 (bottom). Scale bar: 2 $\mu$m. (**D**) Violin plot showing transcriptional silencing induced by Chr 15 expressed Xist transgene as determined by allelic ChrRNA-seq after 24 h Xist RNA induction. The Chr X mESC line without the Xist transgene is shown for comparison. (**E**) As in (D) showing silencing of individual genes across Chr 15 in mESCs expressing Xist transgene on Chr 15 (red circles) compared to parental mESCs (black circles). (**F**) Exemplary widefield images of Xist RNA FISH in Chr X and Chr 15 mESCs after 24 h doxycycline induction. (**G**) Quantitative evaluation of the proportion of mESCs with Xist RNA clusters in Chr X and Chr 15 mESCs after 3 h and 24 h doxycycline induction.

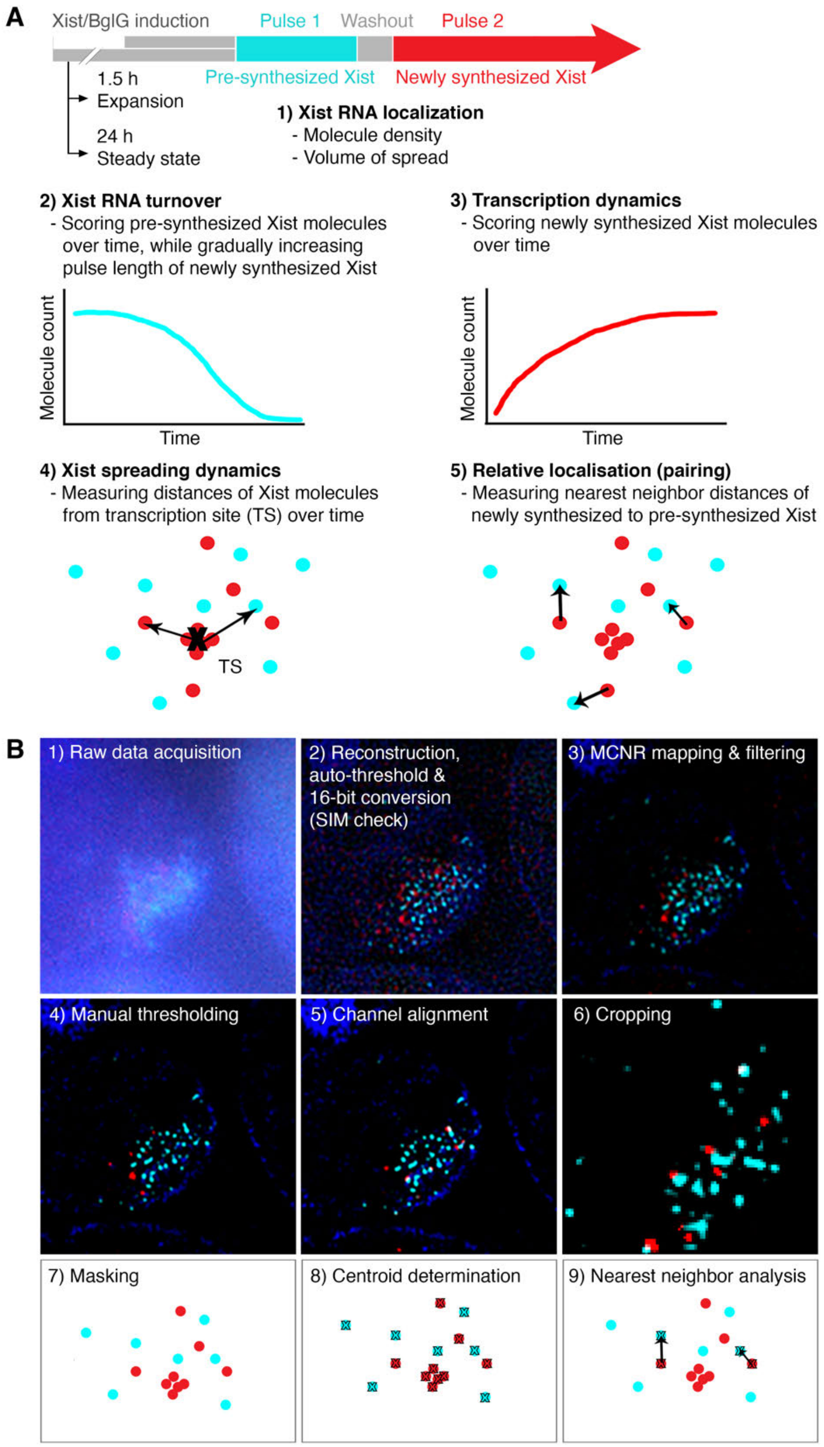


**Fig. S2. RNA-SPLIT image acquisition, processing and analysis workflow.**
(**A**) Schematic illustrating experimental regimens and different applications of RNA-SPLIT. (**B**) Image processing and analysis workflow of RNA-SPLIT samples (see Methods section for details).

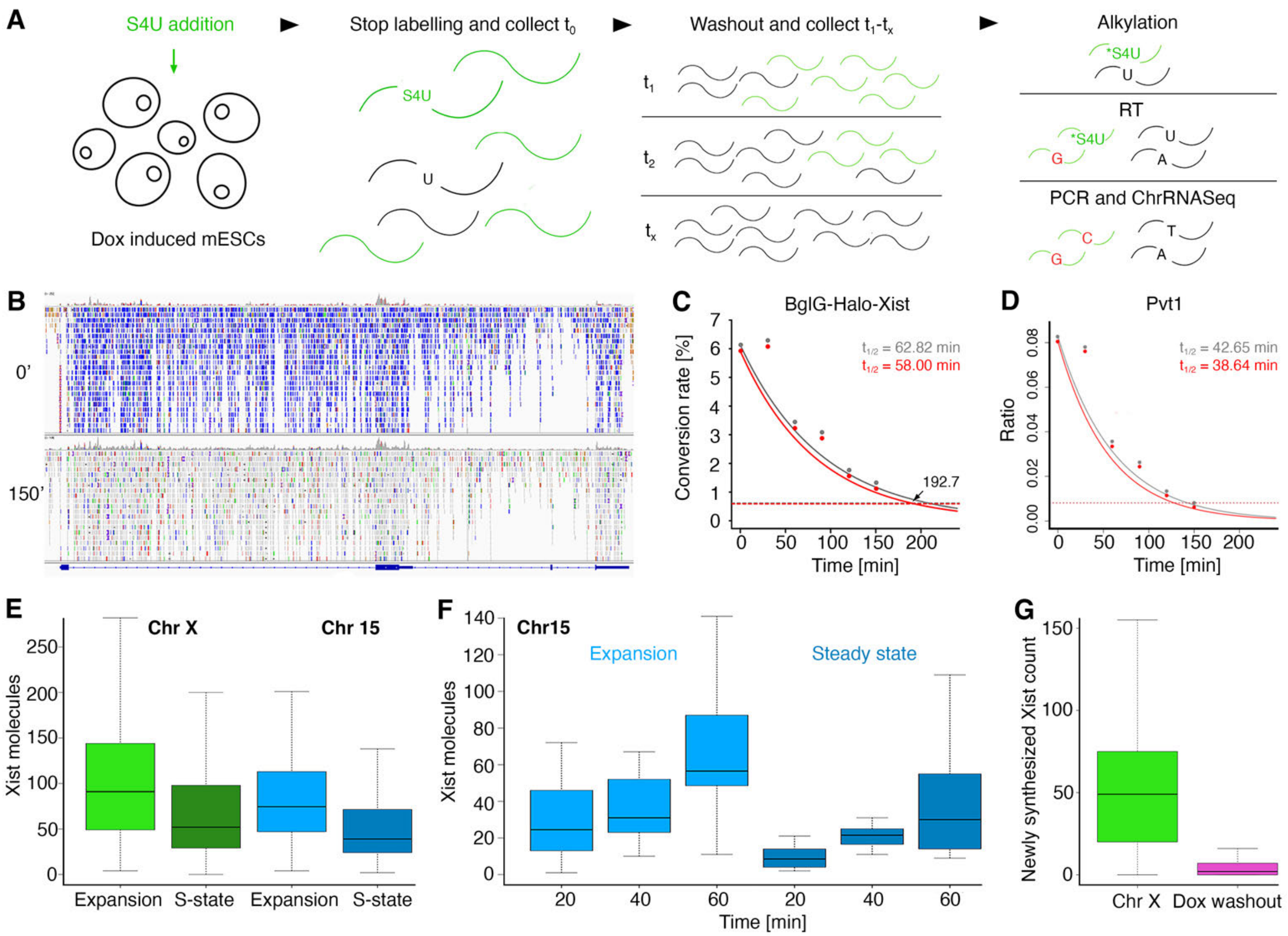


**Fig. S3. Xist RNA stability and rates of transcription are linked.** (**A**) Schematic drawing illustrating the SLAM-seq method for measuring RNA turnover. (**B**) Example showing progressive loss of T to C conversion events (blue) over time (0 min and 150 min) for BglG-Halo-Xist. (**C**) T to C conversion rates over time with fitted exponential decay curves and calculated $t_{1/2}$. Data based on time points at which T to C conversion rates reach at least 10 % of the starting rate. (**D**) T to C conversion rates over time of the lncRNA Pvt1 with fitted exponential decay curves and calculated half-lives. (**E**) Boxplots displaying quantification of Xist RNPs in WT and transgenic line during expansion and steady state. n = 200 cells. (**F**) Boxplots showing Chr 15 Xist transcription rates during expansion and steady state as compared to WT cells, quantified by scoring the number of newly synthesized Xist foci over time. n = 20 cells/time point. (**G**) Boxplots illustrating termination of Chr X Xist transcription (scoring Halo-tagged Xist RNPs in 3D-SIM images) following doxycycline (Dox) washout. n = 100 cells/condition.

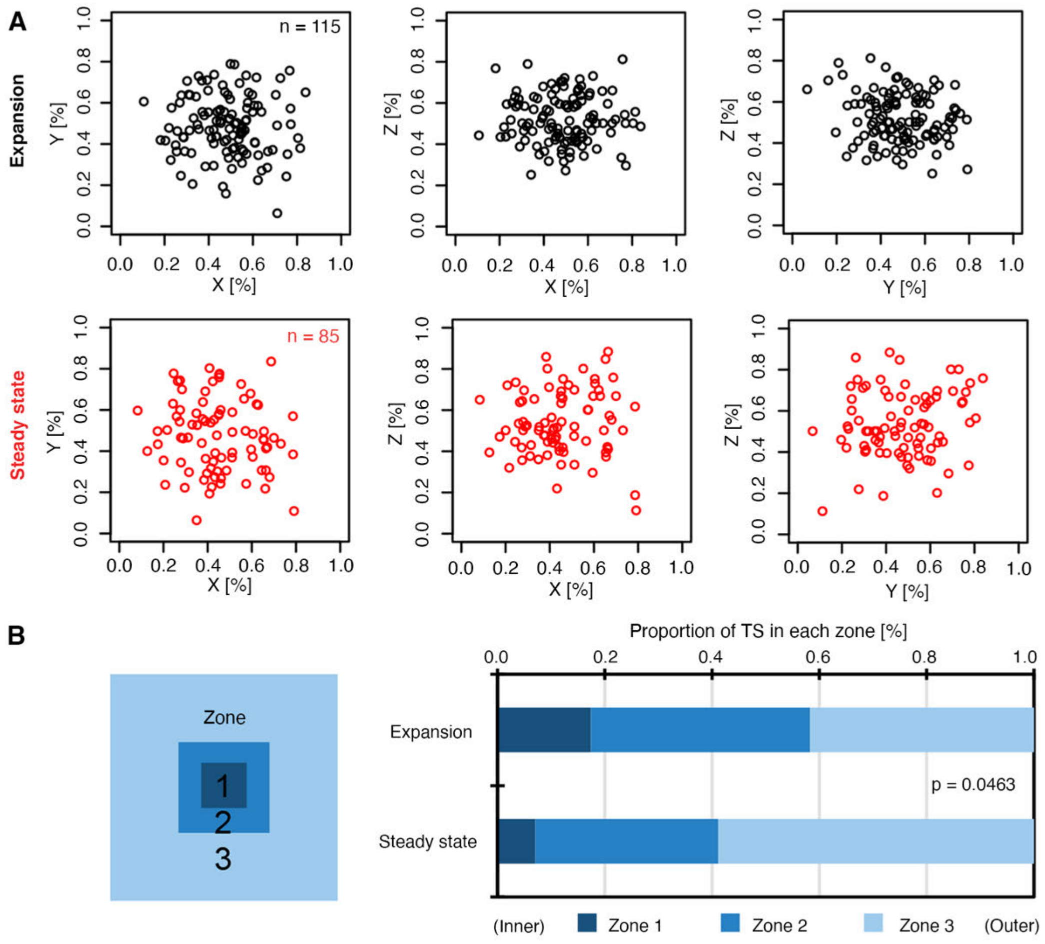


**Fig. S4. Position of the Xist transcription site relative to Xist RNA clusters.**
(**A**) Comparison of Xist transcription site location (normalized to cluster size) in expansion (black) and steady state (red) in 2D from 3 different angles: Y/X, Z/X and Z/Y. (**B**) Proportion of cells with Xist transcription site in zones 1-3 as shown in schematic (left).

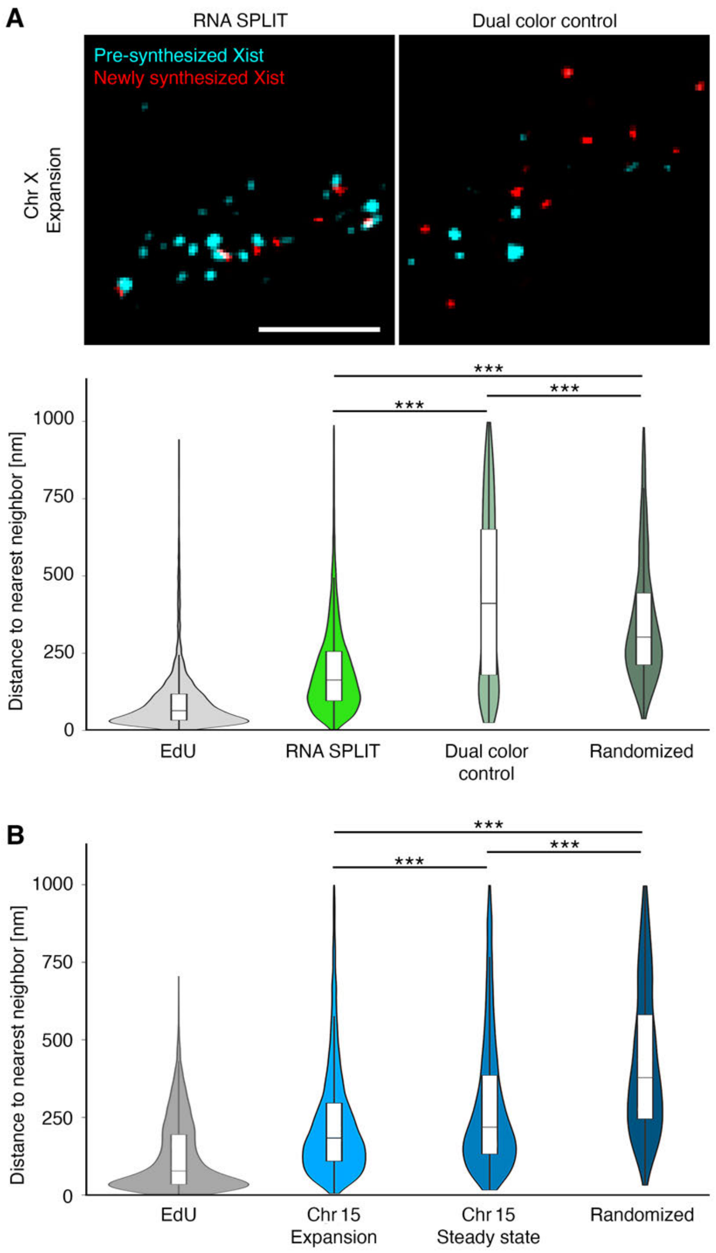


**Fig. S5. Dual-color control to verify Xist RNP coupling.** (**A**) Top: representative 3D-SIM images (single z-section) of RNA-SPLIT of Chr X mESCs with separate addition of fluorophores in expansion phase (left) and dual color control experiment with addition of equal mix of both fluorophores for labelling of pre-synthesized and newly synthesized Xist RNPs. Scale bar: 2 $\mu$m. Bottom: Corresponding violin plots of nearest neighbor distance distributions compared to EdU colocalization and randomized localization controls. n = 120 cells (Chr X expansion) and 20 cells (Chr X dual color control). (**B**) Violin plots as in (A) for Chr 15 Xist transgene mESCs in expansion and steady state phase. n = 200 cells/time point.

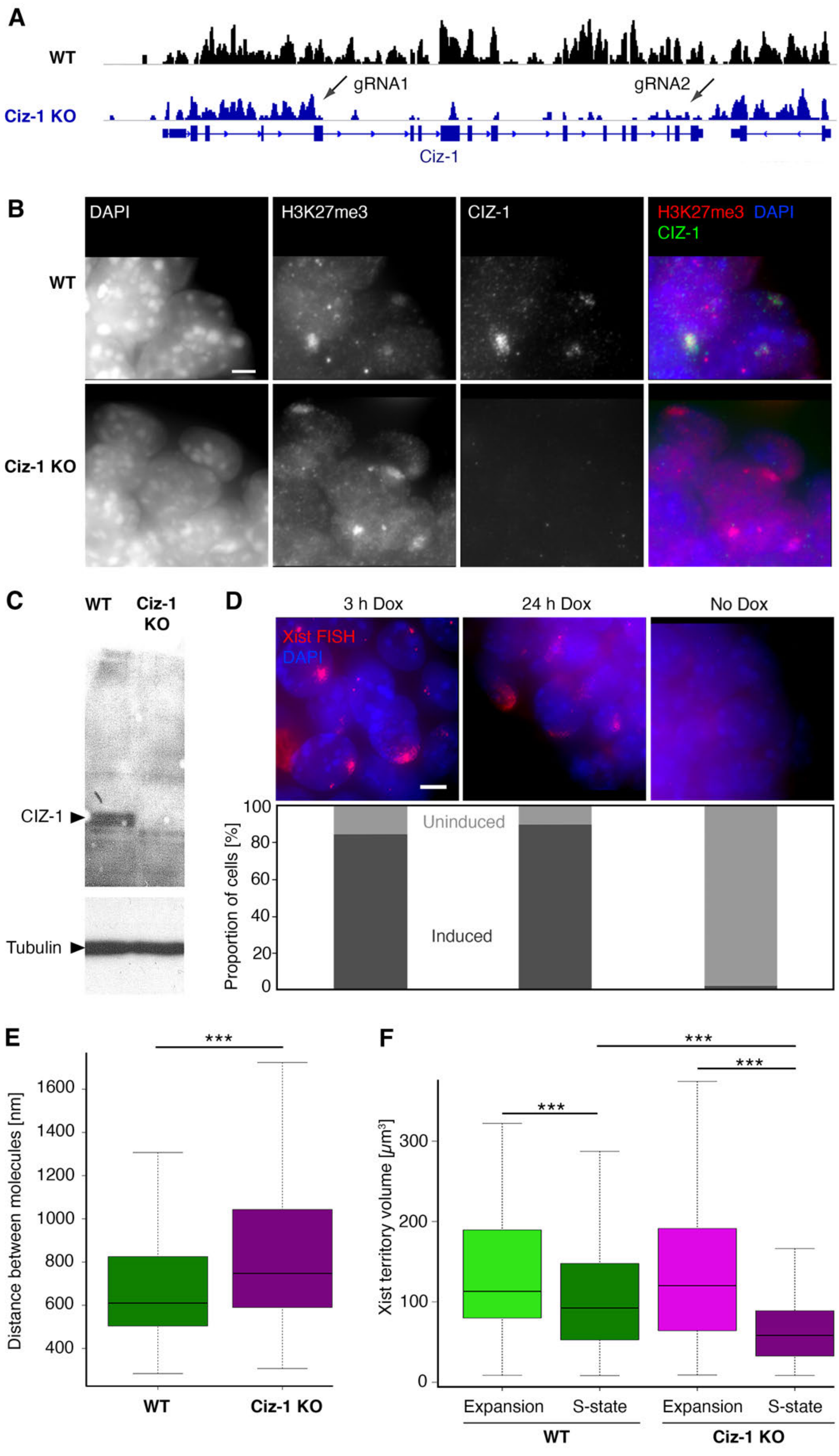


**Fig. S6. Ciz-1 KO mESCs with endogenous inducible BglG-Halo-Xist.** (**A**) UCSC genome browser track showing ChrRNA-seq data confirming CRISPR/Cas9 mediated deletion of a 10.66 kb region of the Ciz-1 locus. sgRNA target sites are indicated (arrows). (**B**)

Representative widefield images (z-projections) of combined immunostaining for H3K27me3 and CIZ-1 in Chr X and Chr X Ciz-1 KO cells 24 h post Xist RNA induction. DAPI counter stain. Scale bars: 5 $\mu$m. (**C**) Western blot for Ciz-1 in WT and Ciz-1 KO mESCs. $\alpha$-tubulin loading control. (**D**) Xist RNA induction in Chr X Ciz-1 KO mESCs 3 h or 24 h after doxycycline addition and no doxycycline control. Top: Representative widefield images (z-projections) show Xist RNA FISH with DAPI counter stain. Scale bar: 5 $\mu$m. Bottom: Quantitative evaluation of induction efficiency. (**E**) Boxplots illustrating quantification of Xist RNP density in Chr X Ciz-1 KO mESCs. n = 240 cells/cell line. (**F**) Boxplots showing Xist cluster volume in Chr X Ciz-1 KO and Chr X mESCs during expansion and steady state phases. n = 200 cells/time point.

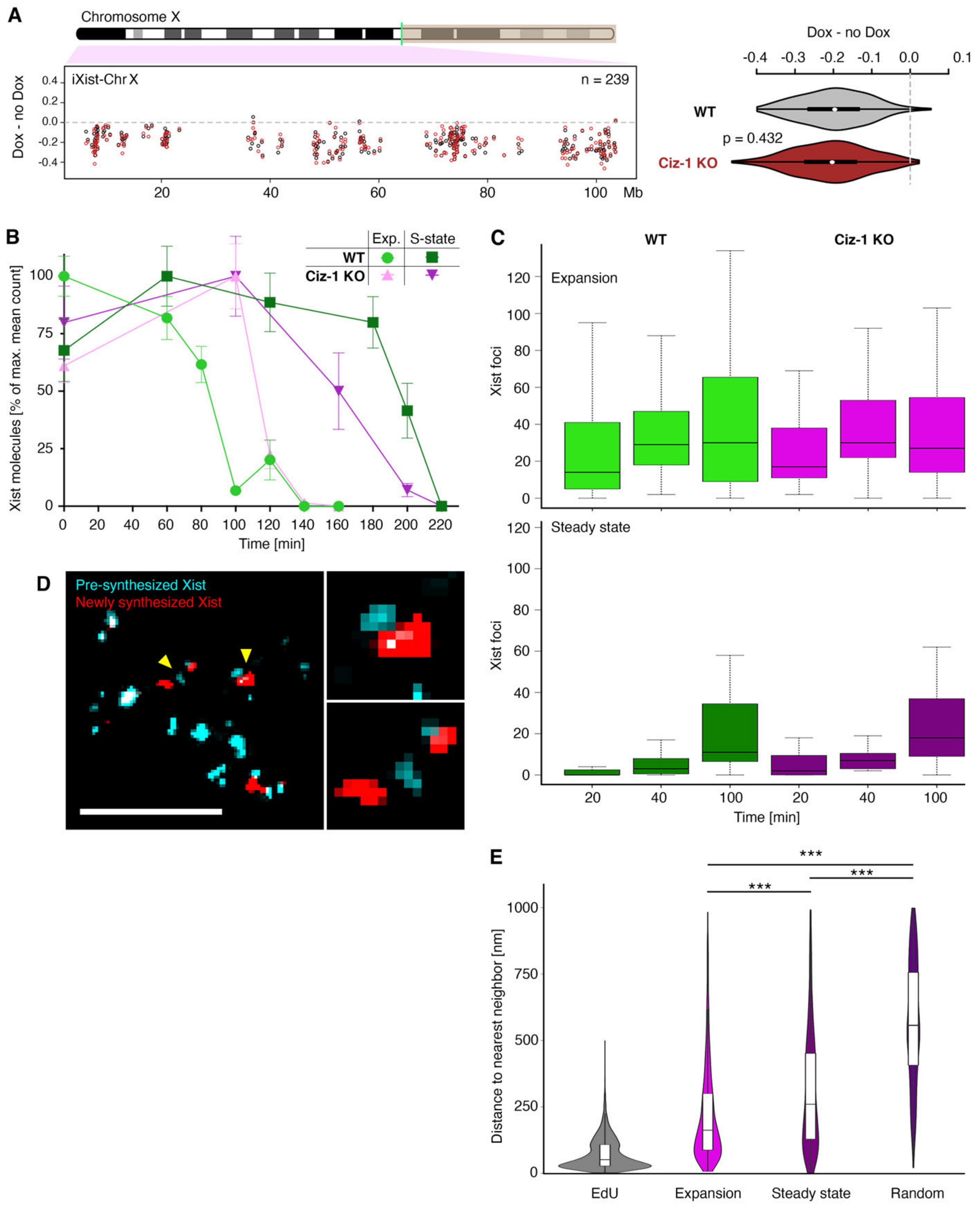


**Fig. S7. Xist-mediated silencing, Xist RNA dynamics and coupling in Chr X Ciz-1 KO mESCs.** (**A**) Allelic ChrRNA-seq analysis showing transcription of individual X-linked genes in Chr X and Chr X Ciz-1 KO mESCs after 24 h of doxycycline induction. Average data from 2 replicates is shown in violin plot (right). Significance determined using unpaired two-sample Wilcoxon test. (**B**) Quantification of Xist RNA turnover in Chr X and Chr X Ciz-1 KO mESCs during expansion and steady state. n = 20 cells/time point. (**C**) Xist RNA transcription rates in Chr X Ciz-1 KO mESCs during expansion and steady state. n = 20 cells/time point. (**D**) Representative 3D-SIM image (single z-section) of RNA-SPLIT experiment using Chr X Ciz-1 KO mESCs with expanded view of selected couplets (arrowheads). Scale bars: 2 $\mu$m (main image) or 200 nm (expanded panels).(**E**) Violin plots showing coupling behavior of Xist RNPs in Chr X Ciz-1 KO cells. n = 200 cells/time point. Significance determined using unpaired two-sample Wilcoxon test.

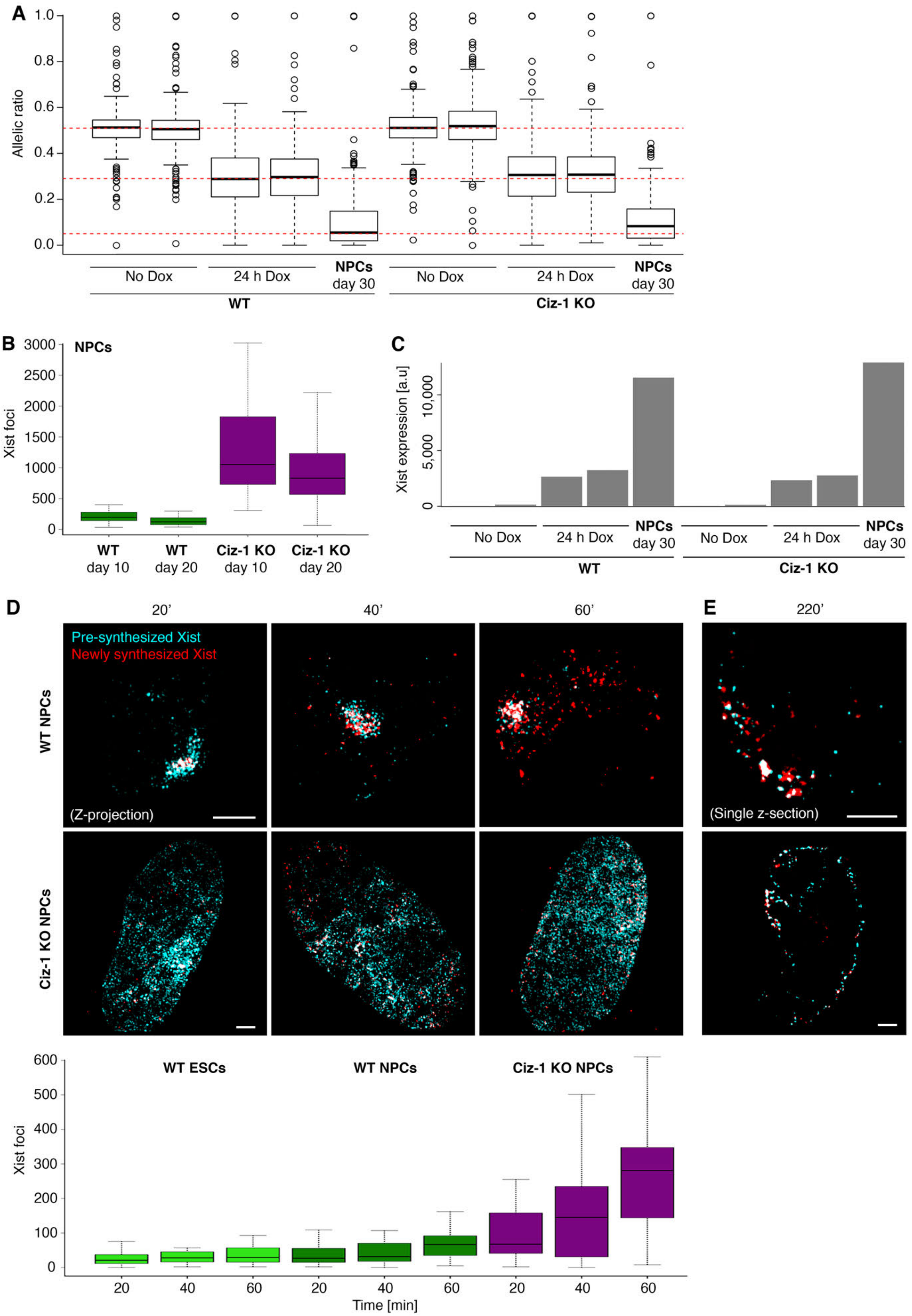
A
Allelic ratio
No Dox
24 h Dox
NPCs day 30
WT
Ciz-1 KO
B
NPCs
Xist foci
WT day 10
WT day 20
Ciz-1 KO day 10
Ciz-1 KO day 20
C
Xist expression [a.u]
D
20'
40'
60'
Pre-synthesized Xist
Newly synthesized Xist
WT NPCs
Ciz-1 KO NPCs
(Z-projection)
E
220'
(Single z-section)
WT ESCs
WT NPCs
Ciz-1 KO NPCs
Xist foci
Time [min]

**Fig. S8. Xist RNA transcription and stability is increased in Ciz-1 KO NPCs.**
(**A**) Boxplots summarizing allelic ChrRNA-seq analysis of X-linked gene expression in Chr X and Chr X Ciz-1 KO mESCs after 24 h of doxycycline induction, and Chr X and Chr X Ciz-1 KO NPCs after 30 days of differentiation. (**B**) Boxplots showing number of Xist RNPs in Chr X and Chr X Ciz-1 KO NPCs after 10 and 20 days of differentiation. n = 200 cells/time point. (**C**) Xist expression levels determined by ChrRNA-seq in Chr X and Chr X Ciz-1 KO ESCs (24 h of doxycycline induction) and NPCs (30 days differentiation). (**D**) Boxplots showing quantification of Xist RNA transcription in Chr X ESCs during expansion, and in Chr X and Chr X Ciz-1 KO NPCs after 20 days of differentiation. n = 20 cells/ time point. Representative 3D-SIM images (z-projections) for Chr X and Chr X Ciz-1 KO NPCs are shown above. Scale bars: 2 $\mu$m. (**E**) Representative 3D-SIM images (single z-section) of RNA-SPLIT experiment illustrating pre-synthesized Xist RNA is present after 220 min in Chr X and Chr X Ciz-1 KO NPCs after 20 days of differentiation. Scale bars: 2 $\mu$m.

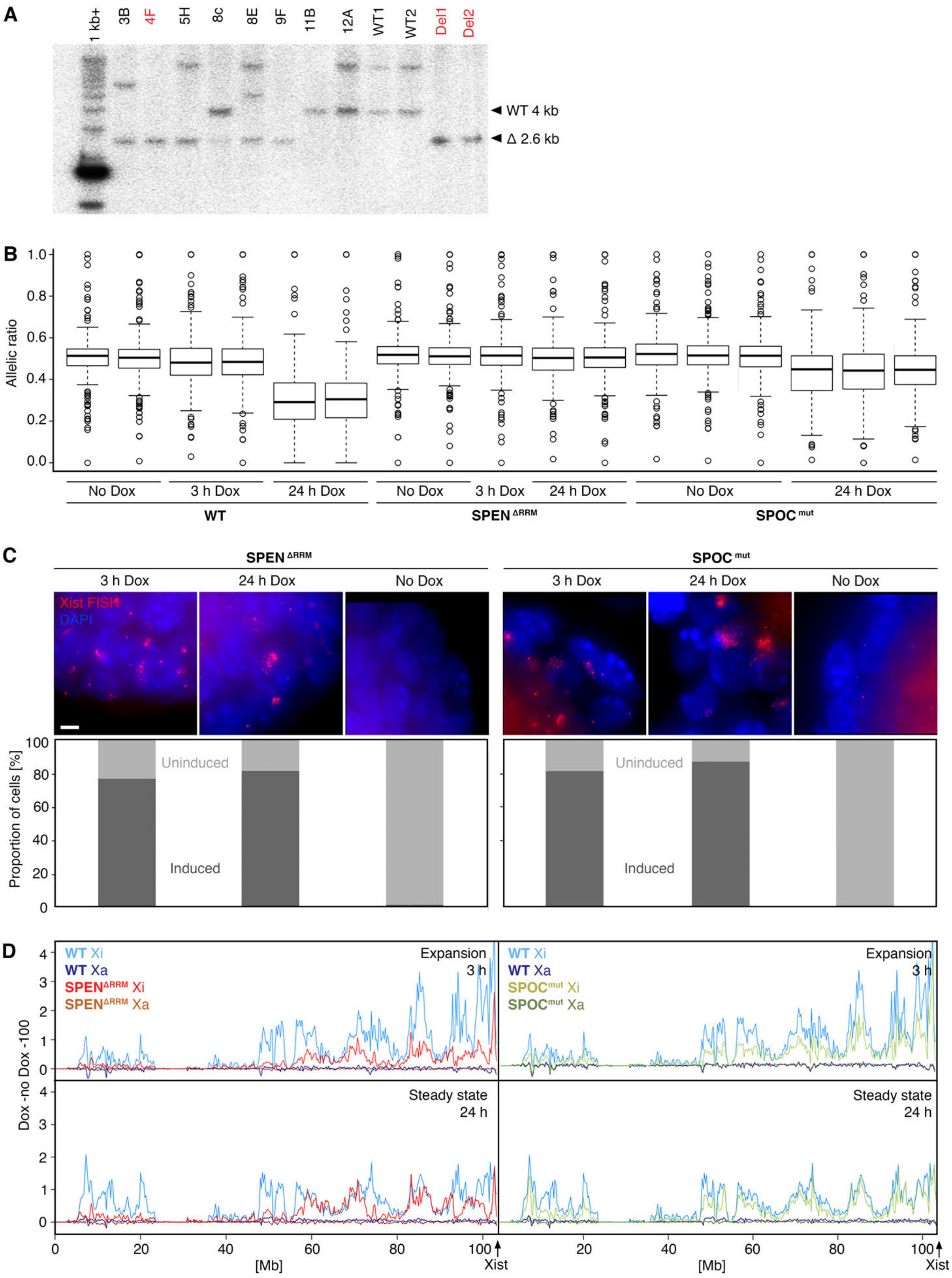
A
1 kb+
3B
4F
5H
8c
8E
9F
11B
12A
WT1
WT2
Del1
Del2
WT 4 kb
Δ 2.6 kb
B
Allelic ratio
No Dox
3 h Dox
24 h Dox
WT
SPEN ΔRRM
SPOC mut
C
Xist FISH
DAPI
Proportion of cells [%]
Uninduced
Induced
D
WT Xi
WT Xa
SPENΔRRM Xi
SPENΔRRM Xa
SPOCmut Xi
SPOCmut Xa
Expansion
3 h
Steady state
24 h
Dox -no Dox -100
[Mb]
Xist

**Fig. S9. Characterization of SPEN RRM del and SPOC mut mESCs. (A**) Southern blot confirming deletion of sequences encoding SPEN RRM 2-4 in Chr X BglG-Halo Xist mESCs. Del1 and Del2 are two previously confirmed SPEN deletions in parental iXist-Chr X mESCs. (**B**) Boxplots summarizing allelic ChrRNA-seq analysis of X-linked genes in Chr X, Chr X SPEN RRM del and Chr X SPOC mut mESCs after 3 h and 24 h of doxycycline induction. (**C**) Top: Representative widefield images (z-projections) of Xist RNA FISH in Chr X SPEN RRM del and SPOC mut mESCs before and 3 h or 24 h after doxycycline induction. Scale bars: 5 $\mu$m. Bottom: Quantitative evaluation of efficiency of Xist induction (scoring presence of Xist clusters). n = 200 cells/time point. **(D**) Plots showing H2AK119ub1 ChIP-seq data (250 kb windows) in Chr X region proximal of the Xist locus split for inactive and active X chromosome alleles in SPEN RRM del cells (left) and SPOC mut cells (right) after 3 h (top) and 24 h (bottom) doxycycline induction. The location of the Xist locus is indicated (arrow).

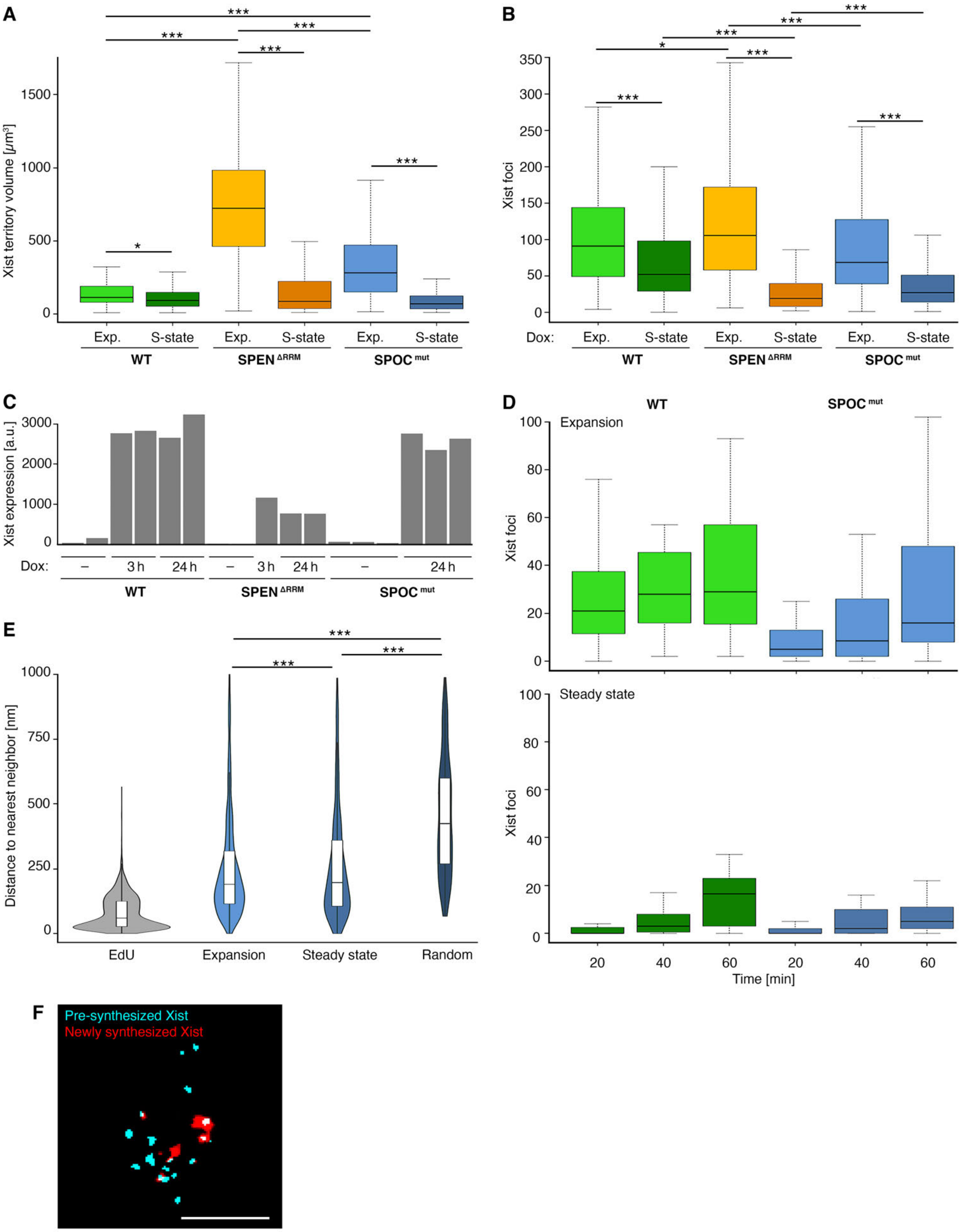

A
B
C
D
E
F
Xist territory volume [μm³]
Xist foci
Exp.
S-state
WT
SPEN ΔRRM
SPOC mut
Dox:
Xist expression [a.u.]
3 h
24 h
Expansion
Steady state
Time [min]
Distance to nearest neighbor [nm]
EdU
Random
Pre-synthesized Xist
Newly synthesized Xist

**Fig. S10. Xist RNP localization, dynamics and coupling in Chr X SPOC mut mESCs.**
(**A**) Boxplots quantifying Xist RNP density in Chr X, Chr X SPEN RRM del and Chr X SPOC mut cells during expansion and steady state. n = 200 cells/time point. Significance determined using unpaired two-sample Wilcoxon test. (**B**) Boxplots quantifying the number of Xist RNPs in Chr X, Chr X SPEN RRM del and Chr X SPOC mut cells during expansion and steady state during expansion and steady state phases. n = 200 cells/time point. Significance determined using unpaired two-sample Wilcoxon test. (**C**) Xist RNA levels (from ChrRNA-seq data) in Chr X, Chr X SPEN RRM del and Chr X SPOC mutant mESCs after 3 h or 24 h of doxycycline induction. (**D**) Boxplots showing Xist transcription rates in Chr X and Chr X SPOC mut mESCs in expansion and steady state phases. n = 20 cells/time point. (**E**) Violin plots showing Xist RNP coupling behavior in Chr X SPOC mut mESCs. n = 200 cells/time point. (**F**) Representative 3D-SIM image (single z-section) showing Xist RNP coupling during expansion phase in Chr X SPOC domain mut mESCs. Scale bar: 2 $\mu$m.

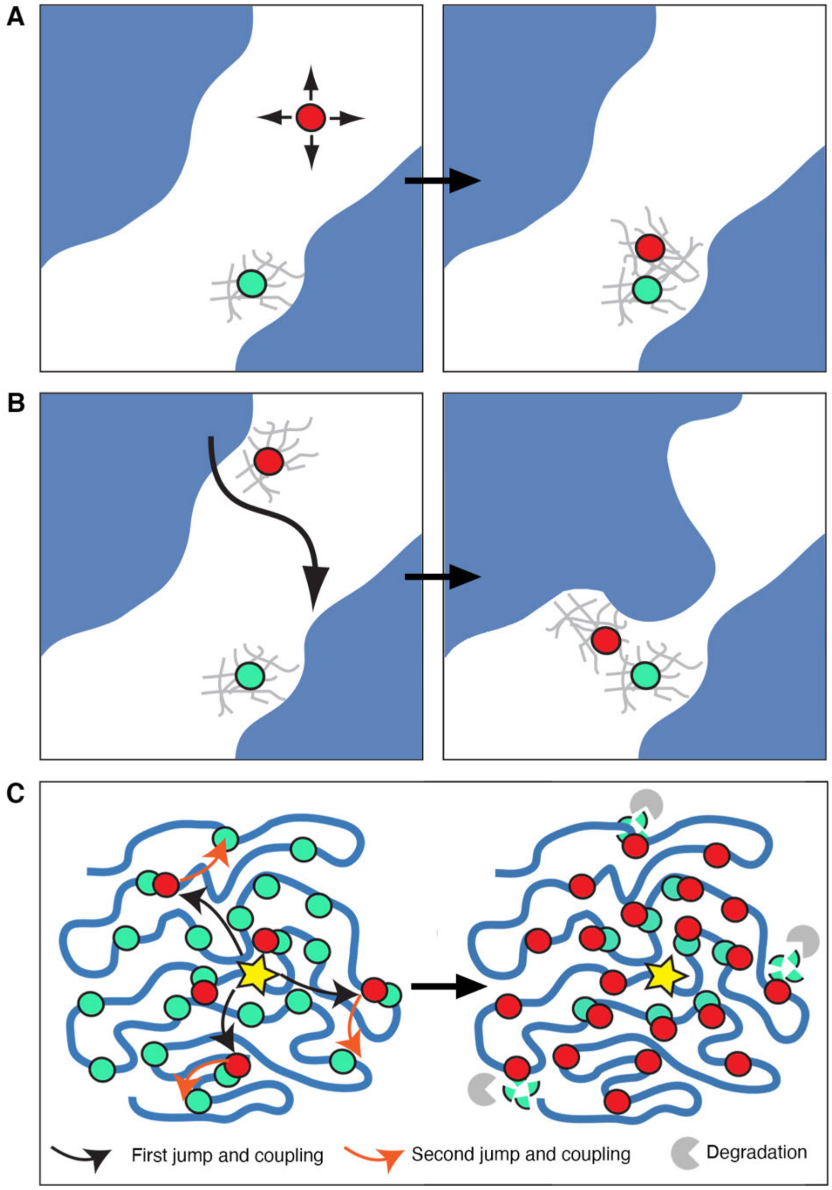


**Fig. S11. Models for coupling behavior and Xist RNP spreading.** (A,B) Schematic illustrating models for Xist RNP coupling. (A) Pre-synthesized Xist RNPs (green) anchor through interaction with nuclear matrix proteins (grey meshwork) and mobile/diffusing newly synthesized Xist RNPs (red, indicated with arrows) show preferential interaction with these pre-established locations. (B) Both pre-synthesized and newly synthesized Xist RNPs are anchored through interaction with nuclear matrix proteins and dynamic movements of chromatin (large arrow) bring together two anchored locations, their association being stabilized by interactions between Xist RNPs and/or the surrounding nuclear matrix. (C) Model for Xist RNP spreading effects during expansion phase seen using RNA-SPLIT. Schematic shows representation of X chromosome at interphase (blue) with Xist transcription site (yellow star), pre-synthesized Xist RNPs (green circles) and newly synthesized Xist RNPs (red circles). Left: pre-synthesized Xist RNPs are shown distributed across the chromosome with newly synthesized Xist RNPs shown as translocating and coupling in a series of jumps from proximal anchor locations close to the

Xist transcription site to more distant locations. Right: illustrating at a later timepoint preferential turnover of pre-synthesized Xist RNPs at distant (peripheral) sites and chromosome wide distribution of newly synthesized Xist RNPs.

**Table S1. Sequences of sgRNAs used for the generation of WT, Ciz-1 KO, SPEN RRM del and SPOC domain mut ESCs, including primers used for PCR screening.**

| Genetic modification | sgRNA 1 oligo | sgRNA 2 oligo | Forward primer 5'→3' | Reverse primer 3'→5' |
|---|---|---|---|---|
| **BgI-stem-loop array** | CATACGTAGT TCCCCGCTCT | N/a | AGTGTGTCTTA CCTATTCCCAT/ GATTCAAGTGG CTCTGAAGTGA | GGTAGGGAGG ATGGCAGTAT |
| **BgIG-Halo** | N/a | N/a | AAAGAGTATGC CTTGCTACCG | TACAGATCCTC AGTGGTTGGC |
| **Ciz-1 KO** | CAGCCTTACA CCACCCCAGA | CAGGGCGTTG CGGGCGTTGA | CTCCGTGCTTT CAATGTGAC | ACTACTCACCT TGCGATTGG/ TACCATCTCCA GATGCTGGG |
| **SPEN RRM del** | GGGGTGTCTC CTGCGCATT | CGGACAAGAC ATTACGATC | GCGCTCCAGCC GAGCCTTCTC | CAGAAAGAGGC GAGGCGTAAAG |
| **SPOC domain mut** | CCCCACTGCG GATCGCCCAG | N/a | GGATCGCCCAG GCCATGGCA/ GGGACACCACA ACGGCCTGTG | CAGCAGCAGGC AGTAGTCGG |

**Table S2: Primary and secondary antibodies used for immunofluorescence labelling.**

| | Primary Antibody | | Secondary Antibody |
|---|---|---|---|
| **H3K27me3** | Anti-mouse, monoclonal, Active Motif (ab61017) | 1:500 | Anti-mouse Alexa Fluor 594, Thermo Fisher Scientific |
| **CIZ-1** | Anti-rabbit, polyclonal, received from the Dawn Coverley lab; purified version used for 3D-SIM | 1:1000 | Anti-rabbit Alexa Fluor 594, Thermo Fisher Scientific |
| **RNA polymerase II** | Anti-mouse, monoclonal, Milipore 05-623 | 1:300 | Anti-mouse Alexa Fluor 594, Thermo Fisher Scientific |
